# Metric and chronological time in human episodic memory

**DOI:** 10.1101/2020.05.11.084202

**Authors:** Hallvard Røe Evensmoen, Lars M. Rimol, Henning Hoel Rise, Tor Ivar Hansen, Hamed Nili, Anderson M. Winkler, Asta Håberg

## Abstract

The relative contributions of metric and chronological time in the encoding of episodic memories are unknown. One hundred one healthy young adults viewed 48 unique episodes of visual events and were later tested on recall of the order of events (chronological time) and the precise timing of events (metric time). The behavioral results show that metric recall accuracy correlates with chronological accuracy for events within episodes, but does not play a role on larger time-scales across episodes. Functional magnetic resonance imaging during encoding and recall showed that metric time was represented in the posterior medial entorhinal cortex, as well as the temporal pole and the cerebellum, whereas chronological time was represented in a widespread brain network including the anterior lateral entorhinal cortex, hippocampus, parahippocampal cortex and the prefrontal cortex. We conclude that metric time has a role in episodic memory on short time-scales and is mainly subserved by medial temporal lobe structures.

---

A primary feature of episodic memory is the preservation of the order of events, also known as “chronological time”^1–3^ (Figure 1e). While mental representation of chronological time preserves the temporal order in which life events occur, it does not itself contain precise information about the timing of events, or “metric time”^3–5^. Indeed, although the mental representation of time clearly is a crucial element of episodic, or autobiographical memory^4, 6–8^, it is still unclear what role representation of metric time plays in the perception of temporal order and in the formation of episodic memories. It is also unclear precisely what regions of the human brain are involved in mental representation of time in episodic memory.

**Figure 1.**
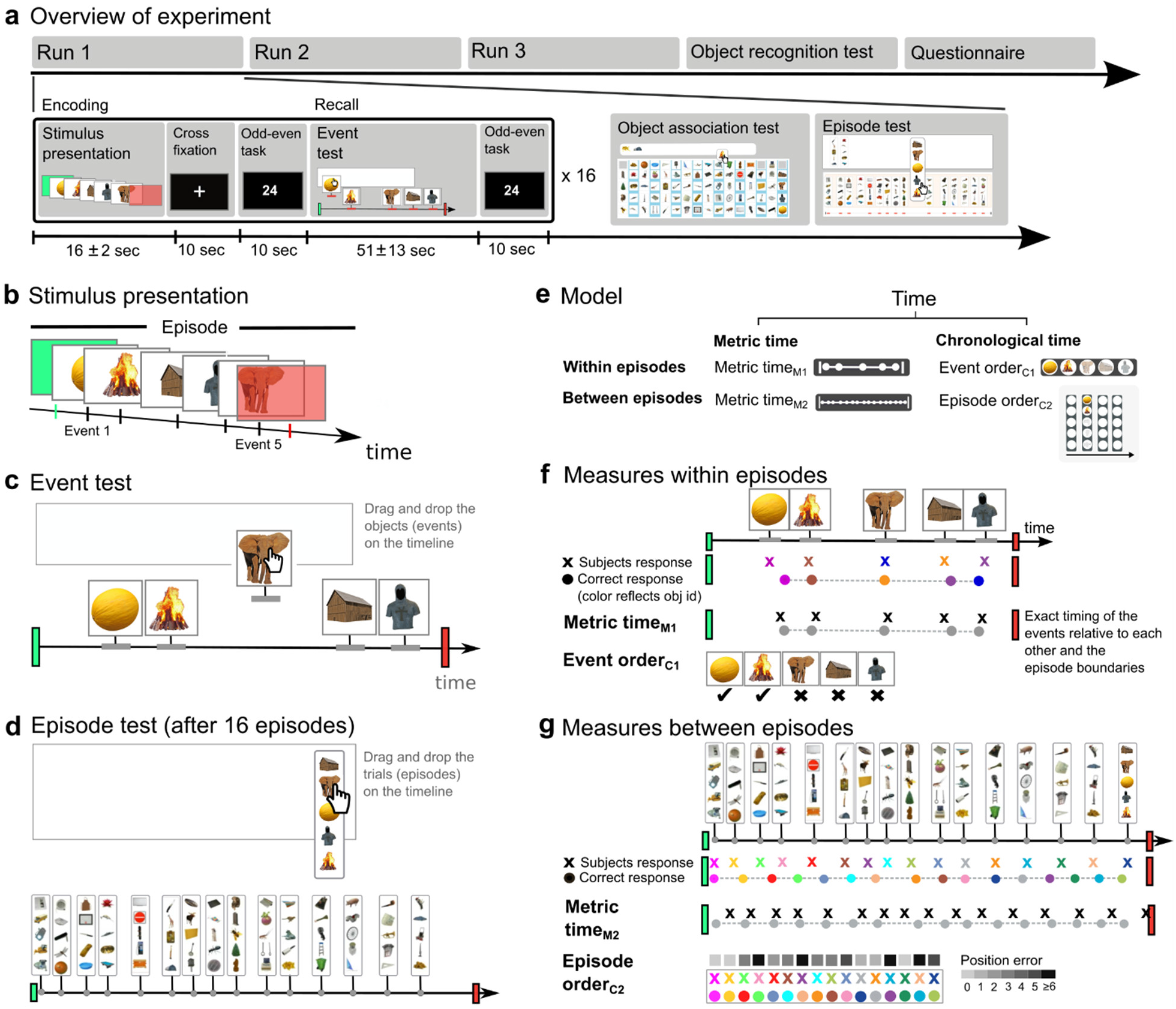
The paradigm and assessment of temporal representation. **a**, Outline of experiment with the top row showing the number of runs and tests performed after all stimuli had been presented (see also Figure S1d), and the bottom row showing the experimental design within a single run. The encoding part of the paradigm involved episodic learning, followed by a crossfixation period, and, subsequently, an odd-even judgment task. There were 16 unique episodes presented within each run. **b**, Each episode consisted of five events (object presentations). A green screen marked the start of the episode and a red screen marked the end (“episode boundaries”). **c**, After the Odd-even judgment, the participant’s recall of the episode was evaluated in the Event test, in which the participant positioned the objects he had seen (i.e., “events”) by dragging and dropping them onto an empty timeline representing the entire episode. **d**, At the end of a run, after having learned 16 event sequences, the participant completed the Episode test and the Object association test (see Figure S1c). In the Episode test, the participant indicated at what point episodes occurred within a run, by dragging-and-dropping the episodes (defined by the objects presented in them) onto an empty timeline representing the entire run. **e**, Representation of time consists of metric time, which represents the exact onset of specific events and episodes, and chronological time which represents the order of the events and episodes. **f**, From the Event test, metric and chronological measures within episodes were obtained. The top row shows the actual response from one participant, with colors indicating object identity and the green and red screen indicating the temporal boundaries of the episode. *Metric time_M1_* reflects the degree to which the participant’s response preserved the exact timing of the events relative to each other (Temporal pattern_M1_) and the exact onset of the events relative to the temporal boundaries of the episode (Temporal boundary_M1_) (see Methods and Figure S1a for a detailed explanation). *Event order_c1_* reflects how many events were recalled in the correct order (independent of Temporal pattern_M1_ and Temporal boundary_M1_). **g**, From the episode test, the participant’s recall of the timing and episode order across episodes was assessed. *Metric time_M2_* reflects whether the participant’s response preserved the timing of all sixteen episodes in a run, relative to each other, as well as relative to the start- and end of the run. *Episode order_c2_* was estimated by calculating, for each episode, how far off the recalled sequence position of the episode (on the timeline) was relative to the correct sequence position.

It is well-documented that the hippocampus and the entorhinal cortex are important for representation of time in rodents^4, 7–9^. Tsao et al. (2018) demonstrated linearly increasing and decreasing electrical neural activity (ramping) associated with temporal landmarks in the anterior-lateral entorhinal cortex (“LEC”; human analogue: alEC^10, 11^) in rats. In the human hippocampus, fMRI activation has been associated with learning temporal sequences of objects^12–14^ and events^15^. However, the distinction between chronological and metric time was not systematically explored in these studies and, thus, any brain activity associated with episodic memory was interpreted as a mental representation of chronological time.

In rodents, neurons that fire at specific timepoints, so-called “time cells,” have been identified in the hippocampus^16, 17^ and the posterior-medial entorhinal cortex (“MEC”; human analogue: pmEC^10, 11^)^18, 19^. However, in these studies, temporal representation on the scale of seconds develops over time and with training. A requirement for a neural code that produces episodic memory is that the mental representation must arise spontaneously and instantaneously (“one-shot”)^3, 8^; it cannot be the result of a learning process and dependent on a training schedule. Therefore, in keeping with the definition of Tsao et al., we focus on investigating whether metric time plays a role in the formation of instantaneous episodic memories.

We asked the following questions: 1) What role, if any, does representation of metric time play in episodic memory across different temporal resolutions or time-scales? 2) Are metric and chronological time associated with the same, or different, regions in the human brain? In particular, are both metric and chronological time represented in the hippocampus and entorhinal cortex, as suggested by previous literature?

We showed human participants sequences of visual events (objects appearing on a screen) and asked them to recall when the events had occurred within each sequence (episode) (Figure 1a). Crucially, the experimental design (Figure 1b-g) allowed us to tease out the separate contributions to episodic memory of metric and chronological time. Our findings demonstrate that (1) representation of metric time has a role in encoding the order of events within an episode (chronology), but does not have a corresponding role on larger time-scales across episodes. Functional neuroimaging data revealed that (2) metric time was represented in the pmEC, as well as the temporal pole and the cerebellum but, perhaps surprisingly, not in the hippocampus. Chronological time was represented in a widespread brain network including the alEC and the hippocampus.

## Results

To explore the relative contributions of metric and chronological time to episodic memory, we exposed 101 right-handed young adults to 48 unique sequences of events (episodes) visually presented on a screen and subsequently evaluated how accurately they were able to recall metric and chronological aspects of the presented sequences. Thus, we obtained measures of metric and chronological accuracy and used these measures to assess whether metric time is encoded at all during episodic memory formation and, additionally, whether encoding of metric and chronological time are related to each other.

Specifically, each episode contained five events occurring in a unique temporal pattern (Figure 1b,c). Each event consisted of an object being presented for 600 ms, giving the participant sufficient time to identify the object^20^. A “temporal pattern” is here defined as the timing of the events relative to each other. The episodes lasted between 8.6 and 22.8 seconds, followed by a 10-second period with a fixation cross on the screen, during which the participants were free to engage in non-stimulus driven encoding^4, 21^. The cross-fixation period was followed by an odd-even judgement task (10 s), the purpose of which was to give the participants time to consolidate the encoded information. Next, the participants were tasked with dragging and dropping the events from each episode onto an empty timeline representing the entire episode (Event test) (Figure 1c). After one run with encoding and recall of 16 episodes, the participants were tasked with recalling which objects had been presented together in different episodes (Object association test) (Figure S1c), and then with dragging-and-dropping the episodes onto a timeline that represented the entire run (Episode test) (Figure 1d). Towards the end of the experiment, the participants were asked to select images of the 248 objects presented during the experiment, among a total of 385 objects (Object recognition test) (Figure S1d). Finally, participants filled in a questionnaire with 22 questions about the strategies they used to successfully encode the event sequences, in order to gauge whether the representation of time was associated with explicit encoding strategies or rather automatic processes beyond conscious control.

To assess the relative contributions of metric and chronological accuracy in the formation of episodic memories, two variables representing metric accuracy and three variables representing chronological accuracy were derived from the participant’s responses to the drag-and-drop tasks (Figure 1e-g). The two metric accuracy variables were (1) Metric time_M1_ for within-episode accuracy and (2) Metric time_M2_ for between-episode (large scale) accuracy. 1) The Metric time_M1_ variable is a combination of two measures: Temporal pattern_M1_, which reflects how accurately the relative timing of the events within an episode is recalled, and Temporal boundary_M1_ which reflects the timing of the events relative to the start- and endpoints of the episode (the green and red screens) (Figure 1f, figure 2, figure S1). We propose that Metric time_M1_ provides a “timestamp” for each event within an episode^3^. 2) Metric time_M2_, reflects whether the participant accurately recalled the timing of all sixteen episodes in a run, relative to each other (Temporal pattern_M2_), as well as relative to the start- and end points of the run (Temporal boundary_M2_)(Figure 1g, figure 2).

**Figure 2.**
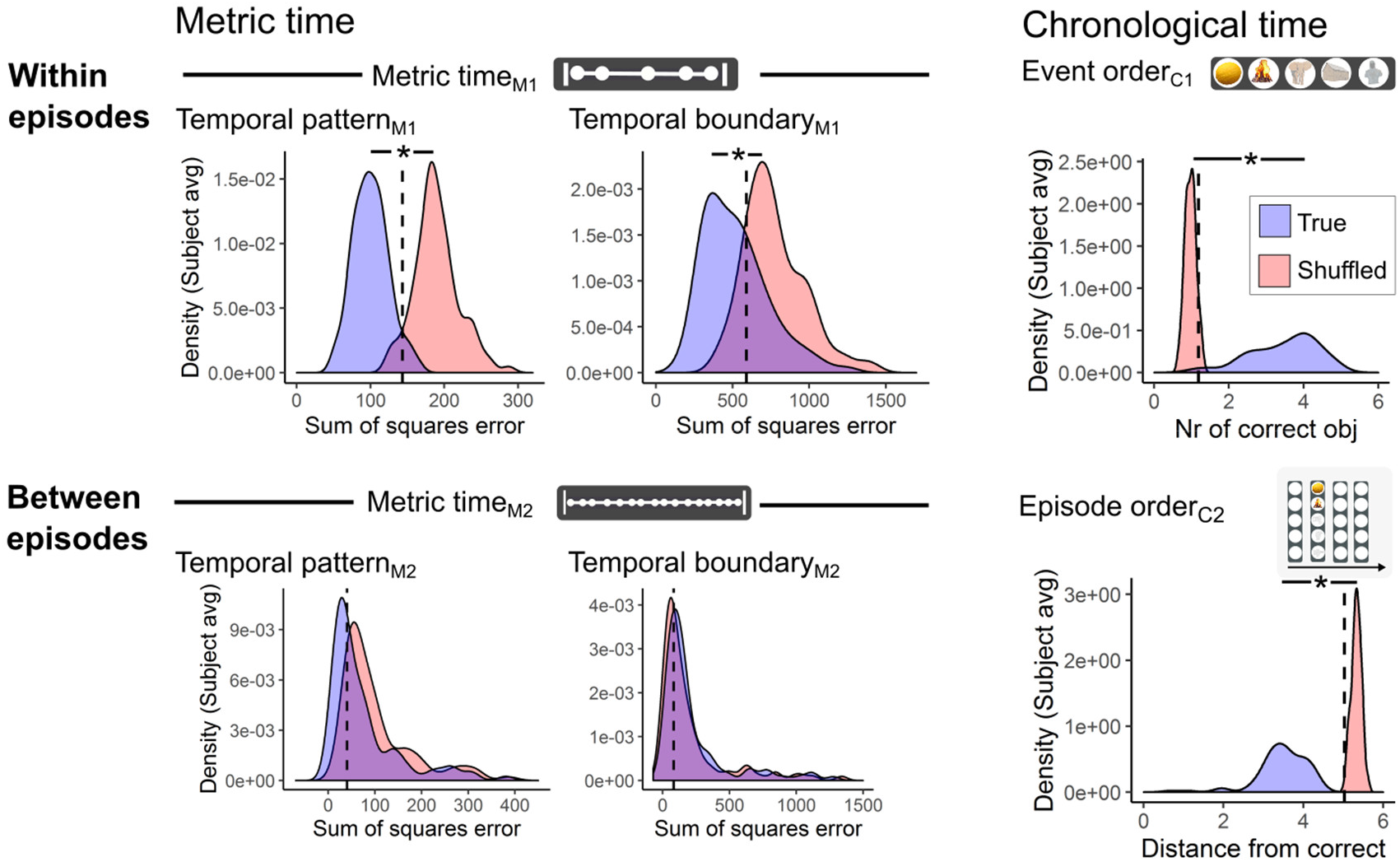
Metric time is only encoded within episodes. The “true” distribution of scores (blue) compared to the “shuffled” distribution of scores (red) (see also Table S1 and Methods). The distributions were based on the average score from each participant. Metric time reflects the degree to which the participant’s response displays the accurate timing between all the events or episodes (temporal pattern) and the accurate timing between the events or episodes and the temporal boundaries (temporal boundary) (see Figure 1 and Figure S1). Recall accuracy of metric time and chronological time were assessed both within episodes (top row) and between episodes (bottom row). The chance level (dotted line) was defined as the cut-off value between the true and the shuffled distribution that showed the most optimal sum of sensitivity (true positive fraction) and specificity (true negative fraction) (see also Figure S2 and Methods). The shuffled distribution of scores was estimated either by comparing the recalled temporal pattern with the correct temporal pattern from all the other episodes (metric measures) (left), or by comparing random sequences of numbers with ordered sequences of numbers (chronological measures) (right) (see Methods). *P < 0.05 (FDR corrected)

The three chronological accuracy variables were (1) Event order_C1_, which reflects how accurately the order of the events within an episode was recalled, (2) Episode order_C2_, which reflects how accurately the order of the various episodes within a run was recalled (Figure 1f). The Episode test was completed three times, once after each of the three runs. Finally, 3) Chronological time_C3_ reflects accuracy of recall for both Event order_C1_ and Episode order_C2_ (Figure 1e). Importantly, the experimental paradigm described here affords exploration of all aspects of temporal representation within the context of episodic memories, in contrast to previous studies which limited the analyses to one, or a few, temporal measures^3, 5^’ ^12–15, 22–24^

### (1.1) Role of metric time: Metric time is only encoded within episodes

To evaluate whether the mental representations of metric time and chronological time were accurate, both within episodes and between episodes, we compared the distribution of the participant’s responses with a shuffled distribution (Figure 2, and see Methods). Chance level was defined as the cut-off between the two distributions when maximizing sensitivity and specificity (Figure 2, Figure S2). For Metric time_M1_ (within episodes) the true and shuffled distributions were clearly segregated (Figure 2 upper panel, Table S1), while for Metric time_M2_ (between all episodes) the two distributions were not significantly different (Figure 2 lower panel, Figure S2c, Table S1). Hence, Metric time_M2_ was not included in mixed linear model analyses evaluating the relationship between metric time and chronological time within-subjects (see 1.2). For chronological time, a clear segregation between the true distribution and the shuffled distribution was observed both within and between episodes, i.e. Event order_c1_ and Episode order_c2_ (Figure 2, Figure S2a, Table S1). This indicates that chronological time (the order in which things happen) is encoded across timescales, whereas exact “timestamps” (metric time) are encoded only within episodes.

### (1.2) Metric and chronological time are correlated within episodes

To test the relationship between metric time and chronological time within-subjects, on a trial-by-trial basis, we employed separate mixed linear models with each of the temporal measures (Metric time_M1_, Event order_C1_, or Episode order_C2_) as response variable. The explanatory variables included the remaining temporal measures, Object association accuracy, Object recognition accuracy, the duration of the episodes, time used on each Event test, differences in strategies during encoding and recall, age, handedness (Edinburgh handedness inventory score), and sex. We found statistically significant associations between Metric time_M1_ and Event order_C1_ (Figure 3, Table S2). Thus, our findings demonstrate that, within episodes, accurate representation of chronological time (Event order_C1_) is dependent on metric accuracy (Metric time_M1_). In addition, metric accuracy was not positively associated with any explicit encoding strategy (Table S2) but rather the result of an automatic, “one-shot” representation.

**Figure 3.**
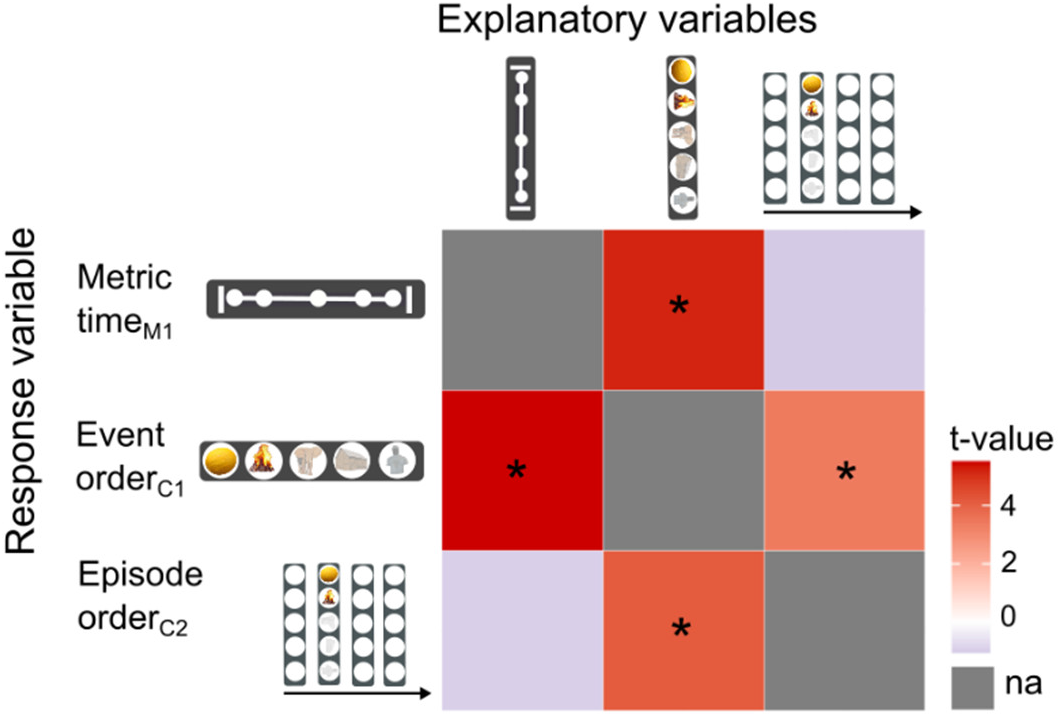
Encoding of metric and chronological time are linked within episodes. Each row defines a behavioral mixed linear model with the row name defining the response variable and the column names the explanatory variables tested for inclusion in the model (see Methods and Table S1). *P < 0.05 (FDR corrected); “na” indicates that the measure did not explain individual variance in the response variable.

### (2.1) Metric time and chronological time are stored in separate neural populations

To evaluate whether metric and chronological time were represented in separate parts of the brain, we investigated whether increased metric accuracy (Metric time_M1_) and chronological accuracy (Event order_C1_ and Episode order_C2_) during encoding and recall of episodes were associated with more dissimilar fMRI activation patterns in the brain. Increased dissimilarity of neural activation patterns are considered to reflect more accurate encoding^13, 25, 26^ due to a reduction in memory interference^4, 26^. We employed a multi-voxel representational similarity analysis (RSA)^27, 28^(Figure S3a). For each voxel in the brain, a 4-mm-radius sphere (“searchlight”) was defined with the target voxel as the center. Every voxel within the sphere had a beta value (from the initial univariate GLM analysis; see Methods) associated with each of the three levels of accuracy (coarse, medium, fine) (Figure S1b). Thus, for each of the variables Metric time_M1_, Event order_C1_, and Episode order_C2_, there was an associated activation pattern (distribution of betas) within the sphere. The RSA analysis tested whether the activation patterns became more or less similar with increasing temporal accuracy (Figure 4a). We considered a brain region to be involved in representation of a specific aspect of temporal processing if the activation pattern dissimilarity in that region was consistently modulated by increasing levels of accuracy (from coarse to medium to fine) both during encoding, including consolidation (stimulus, cross-fixation, or odd-even), and during recall (Event test planning or Event test execution) (see Methods).

**Figure 4.**
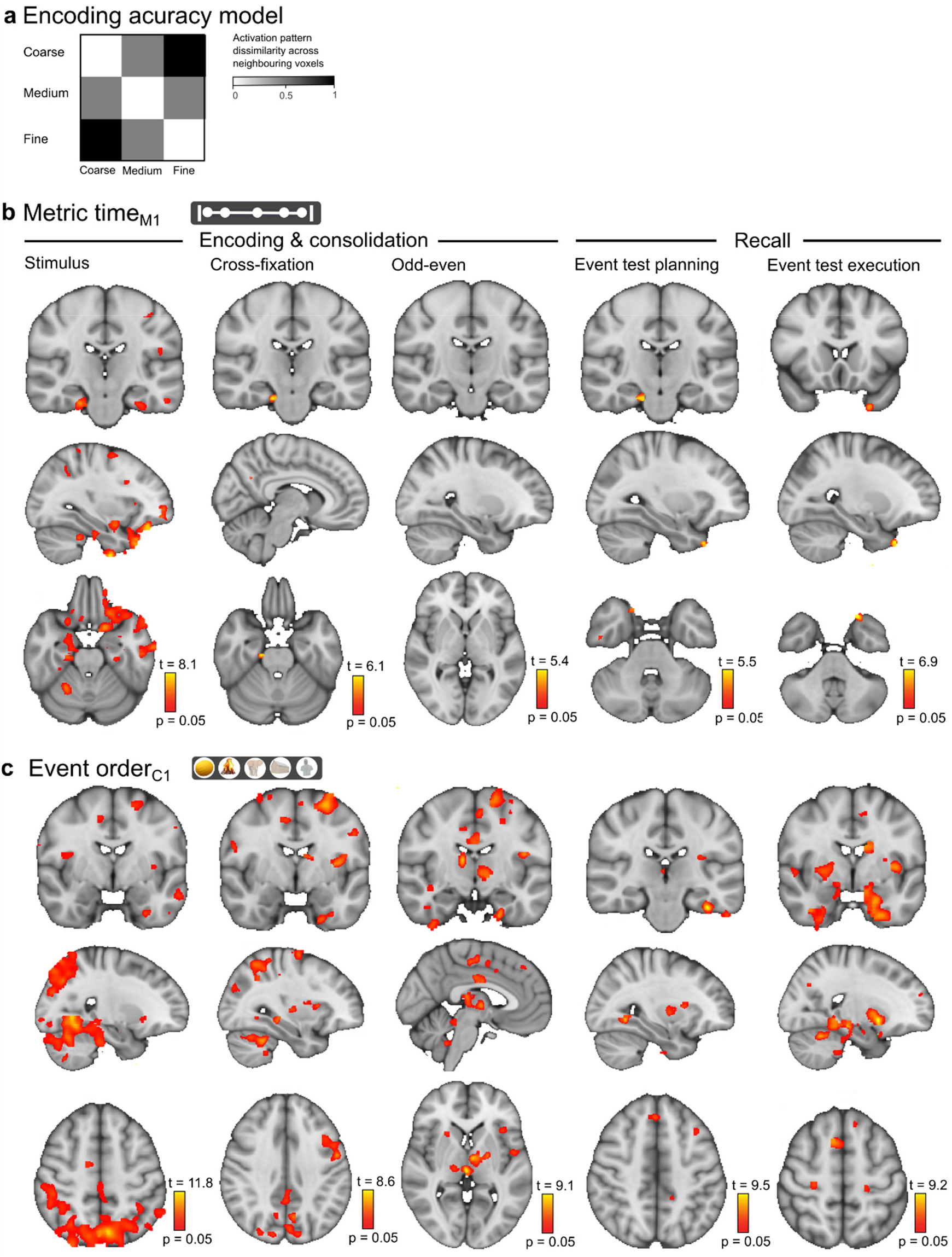
Metric time and chronological time are represented in separate neural populations within episodes. **a**, The statistical model used to test for metric and chronological accuracy. The model predicts a consistent modulation of activation pattern dissimilarity with increasing encoding accuracy (from “coarse” via “medium” to “fine”). **b**, Metric time_M1_. **c**, Event order_C1_ (i.e. chronological time within each episode). Results are shown for encoding which included the Stimulus period (left), Cross-fixation period (mid-left), and Odd-even task (mid), and recall which included the Event test planning (mid-right) and Event test execution period (right) (see Figure 1). p = 0.05 represents the cluster mass corrected thresholds. (For details on activation locations, see Table S3–4).

The RSA analysis revealed that metric and chronological time were represented in separate neural populations within episodes (Figure 4b-c, Table S3–4). Specifically, Metric time_M1_ was represented in a limited network including the pmEc, temporal pole, and cerebellum (Figure 4b, Table S3), while Event order_c1_ was represented in a widespread brain network including the alEc, posterior hippocampus, posterior parahippocampal cortex, perirhinal cortex, temporal pole, posterior prefrontal cortex (inferior-, middle-, and superior frontal gyrus), caudate, putamen, thalamus, insula, cingulate gyrus, temporal gyri, lingual gyrus, fusiform cortex, lateral occipital cortex, visual cortex (V1), precuneus, supramarginal gyrus, angular gyrus, cerebellum, and motor cortices (Figure 4c, Table S4). There were no effects for models where medium accuracy was compared to fine and/or coarse, suggesting that the activation patterns in the brain are consistently modulated by temporal accuracy (coarse-medium-fine). Taken together, these findings show that metric time and chronological time are represented separately in the brain, as predicted by our model (Figure 1e).

### (2.2) Chronological time across timescales is represented in a widespread brain network

Chronological time_C3_, i.e. Event order_C1_ and Episode order_C2_, was represented in the alEc, hippocampus (anterior, intermediate, and posterior), parahippocampal cortex (anterior and posterior), prefrontal cortex (middle frontal and superior frontal gyrus, frontal pole, and orbitofrontal cortex), caudate, putamen, thalamus, insula, cingulate gyrus, superior and middle temporal gyrus, lingual gyrus, temporal occipital fusiform cortex, lateral occipital cortex, visual cortex (V1 and V2), precuneus, supramarginal gyrus, cerebellum, and motor cortices (Figure 5a, Table S5). No independent effect was observed for representation of Episode order_C2_ (Figure 5b, Table S6), suggesting that representation of Episode order_C2_ only becomes relevant when there already is a mental representation of Event order_C1_ (i.e., a representation of the events within the Episode).

**Figure 5.**
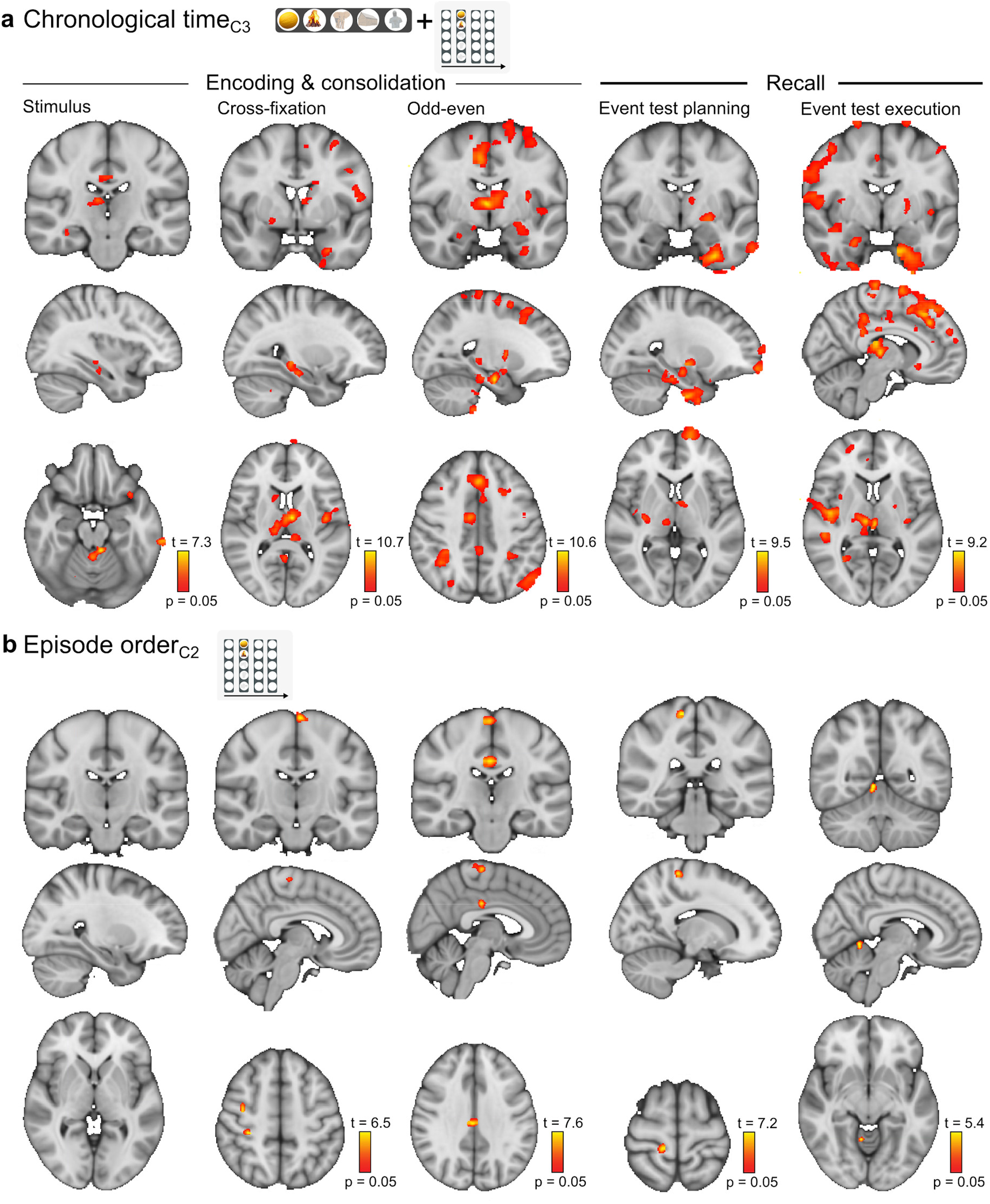
Chronological time across timescales is represented in a widespread brain network. Voxels in the brain that showed consistent modulation of activation pattern dissimilarity as chronological representations became more accurate. **a**, chronological time across timescales. **b**, Episode order (or chronological time between episodes). Results are shown for the Stimulus period (left), cross-fixation period (mid-left), Odd-even task (mid), and the Event test planning (mid-right) and Event test execution period (right). p = 0.05 represents the cluster mass corrected thresholds. (For more details on activation locations, see Table S5–6).

### (2.3) Chronological and metric time are largely represented separately from object identity

To evaluate whether Metric time_M1_, Event order_C1_, and Episode order_C2_ were represented separately from the identities of the objects, we combined the activation pattern dissimilarity analysis with the results from the Object recognition test performed after the MRI scanning. For Object recognition, activation pattern dissimilarity was consistently modulated by increasing levels of accuracy (from 0-3 to 4 to 5 objects recognized) during both encoding and recall in neural populations that to a large extent were separate from those found to be involved in representation of Event order_C1_ and Episode order_C2_. This included the intermediate hippocampus, posterior parahippocampal cortex, putamen, thalamus, insula, posterior cingulate gyrus, temporal gyrus (superior, middle, and inferior), lingual gyrus, occipital fusiform gyrus, lateral occipital cortex, visual cortex V1, occipital pole, precuneus, supramarginal gyrus, cerebellum, and motor cortices (Figure S3b). The unique effect in the occipital fusiform gyri for object recognition is consistent with existing models for how object identity is represented in the brain^29^. Taken together, our results suggest that even though object identity is an essential part of both metric and chronological representation, within the context of episodic memory, object identity is largely represented separately from metric time and chronological time in the human brain.

## Discussion

Our main findings are that metric time is represented in the brain during encoding and recall of episodic memories on a short time-scale (within episodes) and that, contrary to expectation based on previous literature, the hippocampus is not involved in representation of metric time. Rather, metric time is represented in the posterior-medial entorhinal cortex (pmEC), as well as temporal and cerebellar structures, in the human brain. Chronological time is represented in a widespread network that includes the hippocampus and the anterior lateral entorhinal cortex (alEC), as well as the parahippocampal cortex, prefrontal and parietal cortex, the striatum, the cingulate gyrus, insula, and the cerebellum. Within the chronological time network, posterior regions appear to support chronological time on a short time-scale (within episodes), while anterior regions represent chronological time on a larger time-scale (across episodes).

### (1) What role does metric time play in episodic memory?

Our behavioral findings demonstrate that representation of metric time, or precise timing of events within an episode (Metric time_M1_), plays a role in human episodic memory. We found a positive correlation between metric accuracy and chronological accuracy (the order of events), suggesting the brain uses metric time to form an explicit representation of temporal order, perhaps by attaching a “timestamp” to each event (in our experiment, each object presented in an episode). Precise timing is likely to be more important when events are closely spaced as within the current episodes. If these assumptions are correct, representation of metric time is a prerequisite for what Tsao et al. call an automatic “one shot formation of episodic memory”^3^.

Participants recalled metric information above chance level within, but not between, episodes in our experiment. Thus, whereas metric information may be crucial when encoding the order of closely spaced events (within an episode), recall on a larger time-scale appears to be based on a different kind of information. Previous studies suggested that estimates of the duration of intervals between episodes are based on the number of episodes in the intervening interval^14, 24^. In our experiment, the effect of representation of metric time in the brain was larger during the presentation of a sequence of events (the stimulus period) than during recall of the same sequence. Conversely, for chronological time, the effect of representation in the brain increased across the encoding and consolidation periods, and stayed elevated during recall. It was recently proposed that the longer the neural population that represents an episode is active after the episode has ended, the higher the probability that the neural population will integrate information across episodes^4^. To conclude, representation of metric time may primarily be important for memories of individual episodes, and recall on larger time-scales, across episodes, may primarily depend on representation of chronological time.

### (2.1) Are metric and chronological time represented in unique medial temporal lobe regions?

Our brain imaging data show that the posterior-medial entorhinal cortex (pmEC) is involved in representing metric time. This has eluded previous investigations into the role of time in episodic memory, in humans as well as rodents^3, 22, 23, 30^, probably because these previous studies failed to explicitly account for metric time. It is true that the pmEC “time cells” in rats have been found to fire at identical time-points across episodes^18, 19^, but time cell sequences develop with learning across multiple episodes and therefore do not fulfill the requirement for an episodic memory code; that it must arise instantaneously and support memory formation of one-shot experiences^3, 8, 18^. We find that the human pmEC produces instantaneous metric information to support memory formation on a short time-scale. The episodic memories formed in our experiment arose instantly and were largely recalled within 30 seconds, supporting the notion that this is memory formation of “one-shot experiences.” Our findings suggest that a unique metric representation of each episode is engendered by the pmEC in humans, in contrast to the general temporal representation previously observed (across episodes) in the pmEC in rodents^18, 19^.

We demonstrate that the human anterior-lateral entorhinal cortex (alEC) engenders representations of chronological time within and between episodes (across timescales). We found no evidence of metric time representation in the alEC. In rodents, increased electrophysiological ramping activity in the alEC was shown in response to structured behavioral tasks^3^, which was taken to suggest the alEC represents chronological time across timescales^3, 8^.

We find a central role for the human hippocampus in representation of time in episodic memory. This appears to be restricted to chronological time, however, as we find no evidence of hippocampal activation patterns associated with metric time. Conversely, our data demonstrate representation of chronological time across timescales, as well as object identity, in separate neural populations within the hippocampus. This is at odds with the findings of Tsao et al., who report less pronounced evidence for chronological time in the hippocampus and the pmEc, than in the alEc in rodents^3^. It is possible that this discrepancy reflects an inter-species difference between human and rodent brains. Taken together, our findings suggest that the hippocampus represents the chronological “when”, as well as the “what”, component of episodic memory in humans.

### (2.2) How are metric and chronological time represented in the brain beyond the medial temporal lobe?

It has been argued that representation of time in episodic memory most likely depends on a cortico-hippocampal network that “extends far beyond the hippocampus and surrounding structures”^7^. On the other hand, it has also been argued that the representation of time in episodic memory is located in the medial temporal lobe^4, 7–9^. Here, we show for the first time that representation of chronological time is located within a widespread brain network that stretches far beyond the hippocampus and neighboring structures. We find that metric time is represented in a network that includes brain regions not included in existing models for episodic memory, i.e. the temporal pole and the cerebellum (lobule I-IV). These latter brain regions have previously been associated with reproduction of accurate temporal intervals^31^, and the cerebellum is considered one of the core regions within the brain’s “main core timing network”^32^. This suggests a closer link between the brain’s metric timing network and neural networks responsible for the representation of time in episodic memory than has previously been realized.

### (2.3) How are episodic memories integrated on various time scales in the human brain?

We found chronological time on a short scale to be exclusively represented in posterior regions of the hippocampus, the parahippocampal cortex, and the prefrontal cortex. Chronological time across timescales (i.e., across episodes) was represented in anterior regions of the same brain structures. Previous investigations into spatial representation have suggested that the posterior hippocampus subserves finegrained, local representations, whereas the anterior hippocampus subserves coarse, global representations^33, 34^. Furthermore, along the posterior-anterior prefrontal cortical axis there has been shown to be an increase in receptive field size and level of abstraction^35, 36^, consistent with temporal representation from small to large scale.

## Conclusion

Metric time has a role in episodic memory formation and is represented in the pmEC, along with neighboring brain regions and the cerebellum, whereas chronological time is represented in a widespread network including the hippocampus and the alEC. Representation of chronological time on a short timescale (within episodes) involves posterior parts of several brain regions. Anterior parts of the same regions represent chronological time on a larger time-scale (across episodes).

## Methods

### Participants

One hundred and one participants (age: 18 - 33 years, mean 24 years) with no history of neurological disorders, head trauma, previous or current DSM-IV axis I diagnosis of psychiatric illness, including substance abuse, were recruited to the study. All were right handed, as ascertained with the Edinburgh Handedness Inventory^37^, with a mean score of 88.3 ± 13.7 %. Ninety-five participants were male. All participants provided written informed consent prior to participation. The study was approved by the Regional Committee for Medical Research Ethics in central Norway. Twenty-two participants completed the fMRI acquisition (all male), but one was excluded a posteriori because of excessive motion (average frame displacement > 0.3 mm).

### Image acquisition

Functional and anatomical MR images were acquired with a 32-channel Head Matrix Coil on a 3T Siemens Skyra scanner (Siemens AG, Erlangen, Germany). Foam pads were used to minimize head motion. The fMRI stimuli were presented using a LCD monitor with 1280 x 1024 resolution, and the participant using a MRI compatible joystick to make responses (Current Designs, Philadelphia, US). Before the experiment started, the participant was first allowed to familiarize himself with the presentation equipment and the joystick, and then completed practice episodes from the different experimental conditions. Scanning was commenced when complete task compliance was ensured.

T2*-weighted, blood-oxygen-level-dependent sensitive images were acquired during the temporal learning and Event test, using a 2D echo-planar imaging pulse sequence with whole brain coverage. FOV = 220 mm x 220 mm, slice thickness = 3.0 mm (no gap), number of slices = 74, matrix = 74×74 yielding 3.0×3.0×3.0 mm^3^ voxels (TR = 2570 ms, TE = 32 ms, flip angle = 90°). GRAPPA acceleration was used, with a factor of four. The lengths of the functional runs varied between 528 and 781 volumes, due to the variable length of the episodes and the self-paced nature of the recall period. For anatomical reference, a T1 weighted (T1W) 3D volume was acquired using a MPRAGE sequence (TR = 2300 ms, TE = 2.94 ms, FOV = 256 mm x 256 mm x 192 mm, matrix 256×256×192 yielding a resolution of 1.0×1.0×1.0 mm^3^, flip angle = 8°).

### fMRI paradigm

The participants viewed a total of 48 object (event) sequences (episodes) during BOLD fMRI scanning. The large number of episodes increased the likelihood that different levels of accuracy were represented in the participants’ responses. The fMRI paradigm was a block design consisting of three runs, with 16 blocks in each run and one episode per block. Within each block, there were five periods; stimulus presentation (16.0 ± 2.0 sec), cross-fixation (10 sec), an odd-even task (10 sec), an event test (50.5 ± 13.0 sec), and then an odd-even task (10 sec) (Figure 1a). The Event test period was divided into a planning phase and an execution phase. The planning phase was defined as the period until the participant started to move the cursor to place the objects (events) along the timeline, while the Execution phase was defined as the remaining part of the Event test period when the events were placed. Between runs, the participants were given an object association and an Episode test that tested recall of information across the sixteen blocks. Unique event sequences were generated for each participant through random selection from an archive consisting of 480 high quality normative color photographs of objects from a wide range of object categories (https://sites.google.com/site/bosstimuli/). After MRI scanning, the participants were given an object recognition test, a run-order test, and a questionnaire related to strategies used during temporal encoding.

Each episode started with a green screen (0.6 sec), then a random sequence of five unique events was presented for 0.6 seconds, before the episode ended with a red screen (0.6 sec) (Figure 1b). Within each episode, the events were positioned in a unique pattern (Figure 1c). The temporal intervals (durations) between the events and between the events and the start point (green screen) and end point (red screen) were randomly selected with a range between 0.1 – 2 seconds for three of the intervals and a range between 2.1 – 3.5 seconds for the remaining three intervals. This ensured that the length of each episode was within the range of recommended block length for fmri^38^.

The participants were instructed to memorize the sequence in the cross-fixation period, while fixating on a cross on the computer screen. During the odd-even tasks, the participants were instructed to push the right joystick button when an even number (<100) appeared on the screen and the left joystick button when an odd number (<100) appeared (numbers presented at random). The participants were explicitly instructed to focus on getting the odd-even judgments correct, and behavioral data were analyzed to verify compliance. The purpose of the first odd-even period was to separate the stimulus and cross-fixation periods from the Event test, while the purpose of the second odd-even period after the Event test was to provide a clear break between the episodes for the participants, and to function as an implicit baseline for the fMRI data analysis. The web-based paradigm and tests were developed in Meteor (https://www.meteor.com/).

### Behavioral tests between the runs

Between runs, with no ongoing functional image acquisition, the participants were given two tests (1-2) that assessed recall of various temporal and non-temporal information from each of the sixteen recently learned episodes. The participants viewed the tests on the computer screen while lying in the scanner and responded by dragging and dropping events or objects using the joystick.

The Object association test (1) assessed the participant’s ability to recognize which objects belonged together in the same episode (Figure S1c). For each of the sixteen episodes within a run, four of the five objects belonging to the episode were grouped together, and the participant was instructed to select the fifth object among the sixteen missing objects. The participant’s responses were scored as correct or incorrect, according to whether an object was correctly placed

In the Episode test (2), the participant was shown an empty timeline representing the run just encountered. The participant was then instructed to drag and drop each of the 16 episodes onto the timeline in the exact spot corresponding to the episode’s starting point (Figure 1d). From the Episode test, we obtained the Episode order_C2_ and Metric time_M2_ variables (Figure 1g, see below).

### Object recognition test and questionnaire

After MRI scanning, the participants were given an Object recognition test, a Run test, and a questionnaire. In the Object recognition test, the participants were shown the 240 objects from the experiment together with 145 lures. The participants were instructed to click on the objects that they had seen during the experiment (Figure S1d). In the Run test, the participant was shown an empty timeline representing the experiment. The participant was then instructed to drag and drop each of the three runs onto the timeline in the exact spot corresponding to the run’s starting point. The questionnaire was designed to find out which strategies the participants had employed in order to successfully encode the object sequences. The questionnaire had a nine-point scale, ranging from “strongly agree” (9) to “strongly disagree” (1). The participants indicated to what extent they agreed with the following statements: (1) I used rhythm to remember the object sequences; (2) I counted the number of seconds between the objects to remember the duration between the objects; (3) I used memory aids to remember the object sequences; (4) I gave the objects spatial positions; (5) I made up stories to connect the objects; (6) I thought about which categories the objects belonged to; (7) I thought about the names of the objects; (8) I thought about associations I had with the objects; (9) I imagined moving through an environment with the different objects at specific locations; (10) I created subgroups of objects within episodes; (11) I gave each object a number; (12) I continued learning the object sequence during cross fixation; (13) I replayed the object sequence in my mind during cross fixation; (14) I repeated the names of the objects during cross fixation; (15) I compared the most recent object sequence with other object sequences during cross fixation; (16) I started to think about the stimulus period during cross-fixation; (17) I started to think about something else during cross fixation; (18) I relaxed during cross fixation; (19) I continued learning the object sequence during the odd even task; and (20) I focused on the odd and even numbers during the odd-even task; (21) I started to think about something else during the odd-even task; (22) I remember object details.

### Metric measures

Three temporal measures were obtained from the Event test, i.e. Temporal pattern_M1_, Temporal boundary_M1_, and Metric time_M1_. Temporal pattern indicates whether the participant accurately reproduced the relative timing of the five events presented during a given episode, without regard to the identity of the objects presented (Figure S1a). The participant’s response (drag-and-drop onto the 2D overview) was spatially translated such that the geometric center (centroid) of the recalled temporal pattern matched the center of the presented (correct) temporal pattern (as originally presented during the stimulus period), minimizing the root mean square deviation between the patterns^39^. Next, the recalled temporal pattern was scaled (up or down) to minimize the root mean square deviation between the patterns. The purpose of the first transformation (translation) was to disentangle the temporal pattern as such from its relation to the episode’s boundaries (the green and the red square marking the beginning and the end of the episode). The purpose of scaling was to account for the fact that in humans, temporal representations tend to be compressed or expanded^40, 41^. After these transformations, temporal pattern accuracy was obtained as the inverse of the total sum of squares error after all transformations had been performed. That is, for each position (i.e., each event placeholder) in the transformed temporal pattern, the squared error was obtained with respect to the closest position in the correct temporal pattern, and all such squared errors were summed to obtain a total sum of squares error. Temporal pattern was classified as fine, medium, coarse, or “failed.” The thresholds between coarse, medium and fine were defined such that, within participants, the number of episodes in each category was identical (after chance level was determined; see below). Responses were categorized as “failed” if the level of accuracy did not exceed chance level (see below).

The Temporal boundary_M1_ variable reflects how accurately the recalled temporal pattern was placed and scaled relative to the beginning and the end of the episode (Figure S1a). It was defined as the difference between temporal pattern and the inverse of the total sums of squares for the recalled temporal pattern before translation and scaling. Thus, high Temporal boundary_M1_ implies a low degree of translation and scaling relative to the start point (green monitor) and end point (red monitor) of the episode. Responses were classified as fine, medium, coarse, or “failed,” as described above.

The third temporal measure obtained from the Event test was metric time within episodes (Metric time_M1_). Accurate Metric time_M1_ was defined as a response obtaining high scores on both Temporal pattern_M1_ and Temporal boundary_M1_ (Figure S1a). A high Metric time_M1_ score means the participant correctly recalled the temporal pattern as such, i.e. the relative temporal distances between the events, as well as accurately recalling the positioning of this temporal pattern relative to the start and end point of the episode.

From the Episode test, we obtained metric accuracy between episodes (Metric time_M2_) (Figure 1g). Metric time_M2_ reflects the degree to which the participant recalled the exact timing of the episodes relative to each other (Temporal pattern_M2_), and relative to the start- and endpoints of the run (Temporal boundary_M2_) (cf. the detailed explanation of estimation of Temporal pattern_M1_, Temporal boundary_M1_, and Metric time_M1_ above). In addition, because the number of episodes within a run is larger (16) than the number of events within an episode (5), it was pertinent to rule out this as an explanation for the lack of metric time accuracy between episodes. Therefore, we also estimated Temporal pattern_M2_ and Temporal boundary_M2_ for four consecutive episodes (1-4, 5-8, 9-12, and 13-16) within each run (see Figure S2c).

### Chronological measures

Event order_C1_ indicates how accurately the participant recalled the order of the events presented within episodes (Figure 1f). Participant responses were classified into three categories according to the number of events correctly ordered: 2, 3, 4-5 (see below). Chronological time between episodes or Episode order_c2_ was estimated by calculating, for each episode, how far away the recalled episode sequence position was from the correct position in the Episode test (Figure 1g). If the participants for example recalled a specific episode as being the first episode within the run when it was really the third episode, that episode would be given an Episode order score of 2.

### Chance levels

“Failed” was defined as “accuracy (Temporal pattern_M1_, Temporal boundary_M1_, Temporal pattern_M2_, Temporal boundary_M2_, Event order_C1_, Episode order_C2_, Object recognition, Object association) at or below chance level”. For the metric temporal measures, chance level was estimated by comparing the distribution of scores when a recalled temporal pattern was compared to the correct temporal pattern, with the distribution of scores when the recalled temporal pattern was compared to the correct temporal pattern from each of the 48 episodes (excluding the current one) (Figure 2, Figure S2). For the chronological temporal measures, chance level was estimated by comparing the distribution of scores when the recalled event order was compared to the correct event order, with the distribution of scores generated by comparing random sequences of numbers with ordered sequences of numbers 5050 times. For object recognition, we first identified all the objects selected by each participant in the Object recognition test including lures. Then five objects were sampled and the number of non-lures counted. This procedure was repeated 50 times for each participant, and then repeated across all participants to establish the true distribution (for the number of objects correctly recognized on average across episodes). The shuffled distribution was established by sampling five objects from the complete set of objects and lures (originally encountered by each participant) and then counting the number of lures. This was then repeated 5050 times. Chance level for the Object recognition test was subsequently estimated by comparing the true and shuffled distribution. For Object association, chance level was estimated by comparing the distribution of scores when the recalled object associations were compared to the correct object associations, with the distribution of scores generated by comparing random responses (generated using random sequences of 16 numbers) with the correct response (an ordered sequence of the same 16 numbers).

To evaluate whether the true and shuffled distributions were significantly different we used R 3.6.3 (R Core Team (2018). R: A language and environment for statistical computing. R Foundation for Statistical computing, Vienna, Austria. https://www.R-project.org/) and the waddR package (https://github.com/goncalves-lab/waddR). First, the two distributions were normalized by subtracting the mean and dividing by the standard deviation (Z-transformation). Then, the overlap between the two distributions was evaluated using the 2-Wasserstein distance, and 5000 bootstrapped samples combined with a generalized Pareto distribution approximation to get accurate p-values. The significance threshold was corrected for the number of temporal and non-temporal measures tested using a 5% False Discovery Rate (FDR).

The optimal cut-off value between the true and shuffled distributions was estimated using the cutpointr package in R (https://cran.r-project.org/web/packages/cutpointr/vignettes/cutpointr.html). We constructed a receiver operating characteristics (ROC) curve by plotting true positive fraction (sensitivity) vs true negative fraction (specificity) for different thresholds (or cut-off values) between the two distributions (Figure S2). The threshold between the two distributions was estimated by maximizing the sum of sensitivity and specificity from true positives, false positives, true negatives and false negatives for 5000 bootstrapped samples. The optimal threshold was defined as the average across all the samples. For Temporal pattern_M1_ and Temporal boundary_M1_, the thresholds between fine, medium, and coarse episodes were defined so that the average number of episodes in each category was identical, using three significant digits, within participants. For Metric time_M1_, the average number of episodes was 3.4 ± 2.9 for fine, 10.0 ± 5.4 for medium, and 15.8 ± 5.2 for coarse. When measuring accurate recall of event order, only episodes where Temporal pattern_M1_ was above chance level were included. The Event order_c1_ score was divided into correct recall of the sequence positions of four or more events (4-5 events) (avg. no. of episodes: 21.3 ± 13.6), three events (3 events) (avg. no. of episodes: 12.6 ± 6.4), and two events (2 events) (avg. no. of episodes: 6.9 ± 4.9). The Episode order_c2_ score was divided into correct placement within the sequence of 16 episodes with an error of zero to one episodes (0-1 episodes) (avg. no. of episodes: 6.6 ± 4.8), two till three episodes (2-3 episodes) (avg. no. of episodes: 12.0 ± 5.4), or five till six episodes (5-6 episodes) (avg. no. of episodes: 10.9 ± 4.5). For Chronological time, the average number of episodes was 7.6 ± 6.3 for fine, 10.7 ± 4.7 for medium, and 8.2 ± 3.9 for coarse. The Object recognition score was divided into correct recognition of five objects (5 obj) (avg. no. of episodes: 14.6 ± 12.3), four objects (4 obj)(avg. no. of episodes: 15.8 ± 7.1), or zero till three objects (0 – 3 obj)(avg. no. of episodes: 17.6 ± 12.8).

### Behavioral mixed linear models

To evaluate whether there was any association between Metric time_M1_, Event order_C1_, and Episode order_C2_, we employed mixed linear models with maximum likelihood estimates on a trial-by-trial basis. The data was analyzed in R 3.3.2 (R Core Team (2016). R: A language and environment for statistical computing. R Foundation for Statistical Computing, Vienna, Austria. https://www.R-project.org/), using the mixed linear model package lme4 (https://cran.r-project.org/web/packages/lme4/index.html). The R package sjPlot was used for visualization (https://cran.r-project.org/web/packages/sjPlot/index.html). In these analyses, the temporal and non-temporal measures (Metric time_M1_, Temporal pattern_M2_ (for episode 1-4, 5-8, 9-12, and 13-16 within each run), Event order_C1_, Episode order_C2_, Object recognition, and Object association) were employed as response variables in separate models, and explanatory variables were selected on the basis of whether their inclusion improved the AIC (Akaike information criteria) value of the model by using the buildmer package (https://cran.r-project.org/web/packages/buildmer/index.html). In addition, we evaluated absolute measures of goodness-of-fit to determine whether the included variables were indeed informative^42^. We also estimated the variation inflation factors for each model in order to evaluate collinearity between the explanatory variables (using the R package car) (https://cran.r-project.org/web/packages/car/index.html). First, we tested for random intercepts across participants. The fixed effects explanatory variables tested for inclusion in the model were the temporal and non-temporal measures (Metric time_M1_, Temporal pattern_M2_ (for episode 1-4, 5-8, 9-12, and 13-16 within each run), Event order_C1_, Episode order_C2_, Object recognition, and Object association) excluding the measure used as a response variable, when during the experiment the Episodes were presented, duration of the Episodes, time used on each of the Event tests, questions that evaluated strategies used, handedness, age, sex, and finally, a binary variable indicating whether the participant took part in another fMRI study first^33^ and subsequently participated in the behavioral part of this study (outside the MR scanner). The significance threshold was corrected for the total number of explanatory variables, across all models, using a 5% False Discovery Rate (FDR).

### fMRI preprocessing

Preprocessing of the fMRI data was performed using FMRIPREP^43^ (http://fmriprep.readthedocs.io/en/stable/index.html), a Nipype based tool (http://nipype.readthedocs.io/en/latest/). Each T1-weighted volume was corrected for intensity nonuniformity using N4BiasFieldcorrection v2.1.0 and skull-stripped using antsBrainExtraction.sh v2.1.0 (using the OASIS template) (ANTs v2.1.0, http://stnava.github.io/ANTs/). Brain surfaces were reconstructed using recon-all from FreeSurfer v6.0.0 (https://surfer.nmr.mgh.harvard.edu/fswiki), and the brain mask estimated previously was refined with a custom variation of the method to reconcile ANTs-derived and FreeSurfer-derived segmentations of the cortical gray-matter of Mindboggle (http://www.mindboggle.info/). Spatial normalization to the ICBM 152 Nonlinear Asymmetrical template version 2009c (http://www.bic.mni.mcgill.ca/ServicesAtlases/ICBM152NLin2009) was performed through nonlinear registration with the *antsRegistration* tool of ANTs, using brain-extracted versions of both T1-weighted volume and template. Brain tissue segmentation of cerebrospinal fluid, white-matter, and gray-matter was performed on the brain-extracted T1-weighted image using fast (FSL v5.0.9). Functional data was motion corrected using *mcflirt* (FSL). This was followed by co-registration to the corresponding T1-weighted image using boundary-based registration with 9 degrees of freedom, using bbregister (FreeSurfer v6.0.0). Motion correcting transformations, BOLD-to-T1w transformation and T1w-to-template (MNI) warp were concatenated and applied in a single step using *antsApplyTransforms* with Lanczos interpolation.

### Activation pattern similarity analyses

The fMRI data were first subjected to a univariate (single-subject) analysis in SPM 12 (https://www.fil.ion.ucl.ac.uk/spm/software/spm12/). Brain activity was modeled using a general linear model (GLM) with the default options. Two models were used to investigate associations between accuracy measurements and activation pattern dissimilarity. The first model was used to investigate Metric time_M1_, Event order_C1_, Episode order_C2_, Chronological time_C3_, and the second was used to investigate Object recognition. In the first model, the explanatory variables were, in addition to intercept, the temporal measure (with four levels: fine, medium, coarse, and failed) for the stimulus period, the cross-fixation period, the odd-even period that followed the cross-fixation period, and the Event test planning and execution period resulting in a total of 21 regressors (see Figure S3a). The odd-even period after the Event test served as an implicit baseline. In the second model, the levels of accuracy for Object recognition were 0-3 (coarse), 4 (medium), and 5 (fine); here there was no “failed” condition, resulting in a total of 16 regressors for Object recognition. These models allow the BOLD response in a given voxel to be non-linearly modulated by the respective level of accuracy, and do not assume a similar spatial pattern for each of these levels, which then becomes relevant for the RSA analysis (described below). One participant was excluded from the event order analysis, one from the Chronological time analysis, and two from the Object recognition analysis due to lack of episodes in one level of encoding. The effect of timepoints associated with abnormal shifts in signal change due to head motion or severe artifacts were removed from the analysis, by estimating the root mean square variance over voxels (DVARS) and including a regressor for each timepoint with an abnormal DVARS value (https://fsl.fmrib.ox.ac.uk/fsl/fslwiki/FSLMotionOutliers)^44^.

In order to test for associations between encoding accuracy and changes in activation patterns, we used a multivariate representational similarity analysis^27, 45^, implemented in MATLAB (R2018a, Mathworks, Natick, Massachusetts, U.S.A.). The multivariate analysis was restricted to a gray matter mask, in order to reduce noise and improve classification accuracy^46, 47^. The gray matter mask was based on the Harvard Oxford Structural Atlases and the MNIfnirt cerebellar atlas (probability threshold set to 50 percent) (part of FSL; http://fsl.fmrib.ox.ac.uk/fsl/fslwiki/Atlases). A spherical “searchlight” was obtained for each voxel using a maximum radius of 10 mm to create searchlights consisting of 10 voxels (each 3 mm^3^). This resulted in an average searchlight radius of 4 mm, previously shown to be optimal for detection performance^46^. Non-spherical searchlights were allowed for voxels close to the borders of the grey matter mask to make sure that each searchlight contained the same number of voxels. The searchlights were investigated separately and independently of each other. For each searchlight, three activation maps were generated, containing the betas from the univariate GLM analysis for “fine”, “medium”, or “coarse” (2, 3, 4-5 for Event order, 1-2, 3-4, and 5-6 for Episode order, and 2-3, 4, 5 for Object recognition) with odd even as implicit baseline (see Figure S3a). A multivariate noise normalization was then applied to these activation maps, by first extracting the GLM residual for each voxel, then using those residuals to create a covariance matrix between all voxels, and finally using that covariance matrix to perform a spatial pre-whitening of the regression coefficients^28^. The next step was to generate pairwise activation pattern dissimilarity maps for fine vs. medium, fine vs. coarse, and medium vs. coarse, using an Euclidean distance measure (equivalent to computing the Mahalanobis distance between activity patterns)^28^. Subsequently, in order to assess whether the activation pattern dissimilarity was consistently modulated across all levels of accuracy, the dissimilarity maps were correlated with a model representing an increasing order of dissimilarity from coarse through medium to fine (Figure 4a, figure S3a). Thus, for the central voxel of the searchlight region a correlation score was obtained that expressed the degree to which the similarity of activation patterns centered around that voxel were consistently modulated by accuracy (coarse-medium-fine). Kendall’s tau_A_ was used to assess whether the observed activation pattern dissimilarities could be predicted by the accuracy model, because it also allows monotonic (not strictly linear) relationships (Nili et al., 2014). Smoothing was performed on the single-subject level after estimation of the activation pattern dissimilarities, so as not to reduce the “spatial fine structure of the data”^46^, and the resulting correlation maps were smoothed with a Gaussian kernel of 6 mm. To test whether the correlation effects were significantly different from zero at a group level, the smoothed correlation maps were tested non-parametrically with one-sample t-tests using the program Randomise^48^, part of the FSL software package. Inference used cluster mass statistic^49^, with a cluster forming threshold set at p=0.0001. Clusters were considered significant at p=0.05, corrected for multiple testing using the non-parametric distribution of the maximum statistic. For visualization purposes, the data were resampled to 1mm^3^ resolution.

### Data and software availability

Due to privacy concerns and official regulations, the ethical and governance approvals for this study do not permit the MRI data to be made available in a public repository. Data in this manuscript can be accessed by qualified investigators after ethical and scientific review (to ensure the data is being requested for valid scientific research) and must comply with the European Union General Data Protection Regulations (GDPR), Norwegian laws and regulations, and NTNU regulations. The completion of a material transfer agreement (MTA) signed by an institutional official will be required. The software developed at NTNU cannot be made freely downloadable according to NTNUs regulations on innovations (patented and not-patented) but can be made available upon formal agreements between academic institutions.

## Acknowledgement

We thank Linnea Marie Dramdal Borg for helping with the data collection, Øyvind Salvesen for valuable discussions related to statistical analysis of the behavioral data, Kam Sripada for valuable input to the manuscript, and the staff at the Department of Medical Imaging at St. Olavs Hospital in Trondheim for assistance with imaging protocols and data acquisition. This work was supported by the National advisory unit for fMRI in Norway and the Department of Neuromedicine and Movement Science, NTNU.

## Author contributions

H.R.E. designed the experiment, conducted the experiments, analyzed the data, and wrote the paper; L.M.R. contributed to the design of the experiment, data collection, analysis of the data, and the writing of the paper; H.H.R. contributed to the design of the experiment, data collection, analysis of the data, and the writing of the paper; T.I.H contributed to the design of the experiment and the writing of the paper; H.N. contributed to analysis of the data and the writing of the paper; A.W. contributed to analysis of the data and the writing of the paper; A.H. contributed to the design of the experiment and the writing of the paper.

## Supplementary information

**Figure S1.**
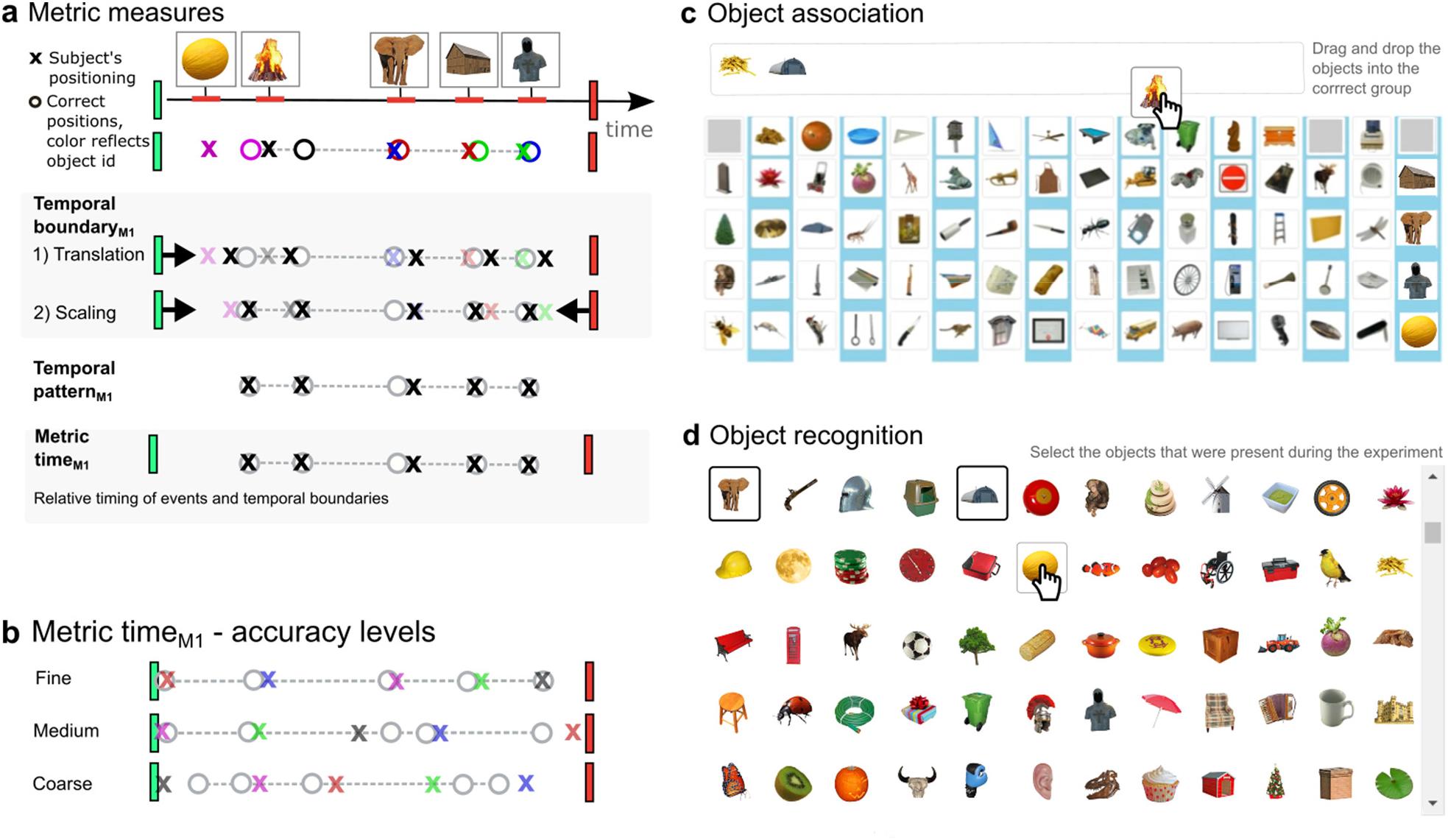
Metric and non-temporal measures. **A**, *Temporal boundary_M1_* reflects the degree to which the temporal pattern, as recalled by the participant, had to be translated and scaled in order to perfectly align with the start and the end of the episode. High Temporal boundary implies low degree of translation and scaling. Correction for scaling was applied because temporal representations are often compressed or expanded^40,41^. *Temporal pattern_M1_* reflects how accurate the timing of the events (objects) relative to each was recalled, after correction for scaling and translation relative to the temporal boundary. Both temporal boundary and temporal pattern are independent of event identity. *Metric time_M1_* reflects the degree to which the participant’s response preserved temporal pattern and temporal boundary. **B**, Levels of recall accuracy for Metric time_M1_, illustrated using actual responses from the participants. **c**, After sixteen event sequences the participants were tasked with recalling which objects had been presented together (*Object association test*). **d**, Towards the end of the experiment the participants were tasked with selecting the objects that had been part of the experiment (*Object recognition test*).

**Figure S2.**
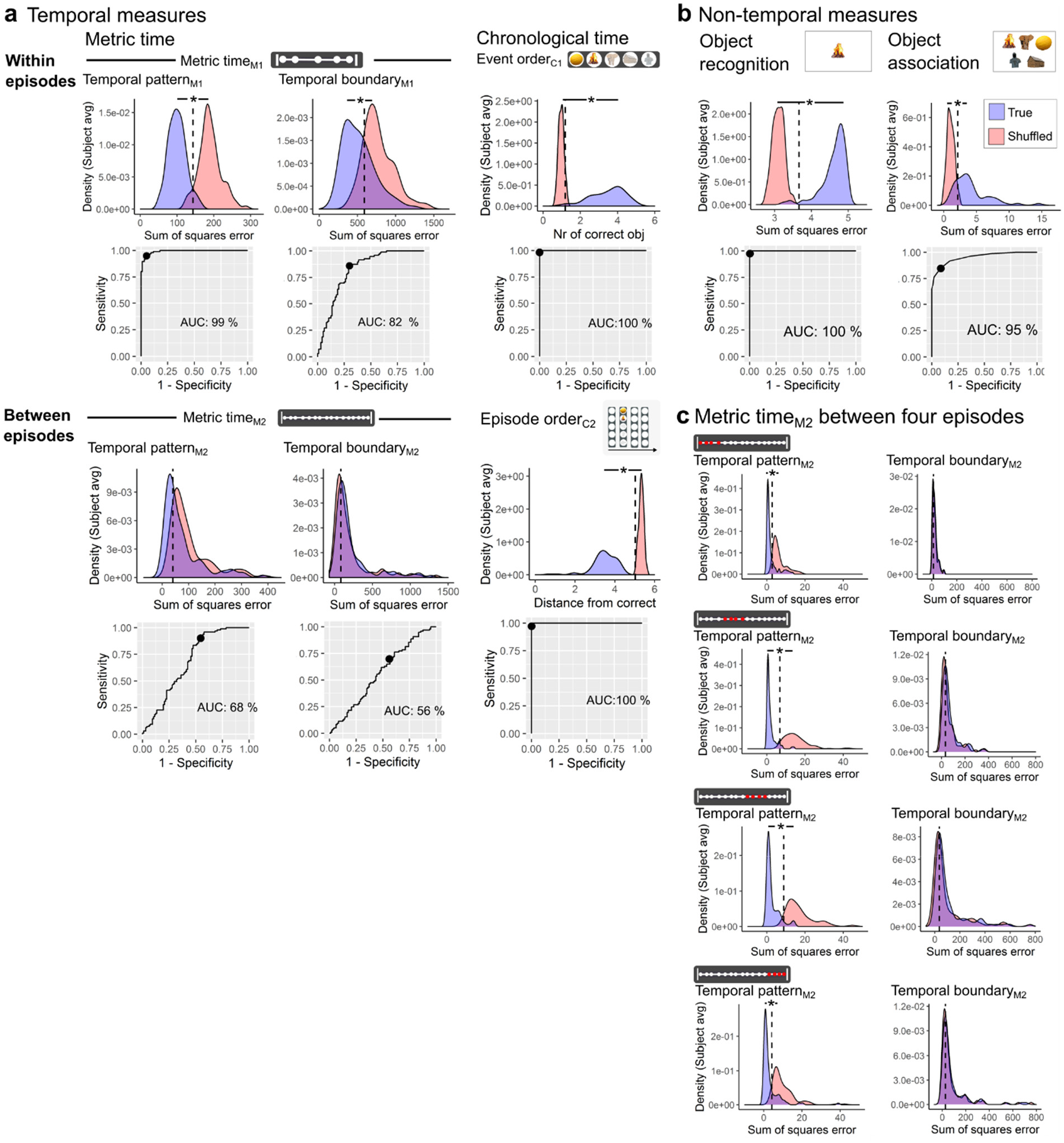
The encoding of temporal and non-temporal aspects of episodic memories. The “true” distribution of scores (blue) compared to the “shuffled” distribution of scores (red). The distributions were based on the average score from each participant. Receiver operating characteristics (ROC) curve plots show true positive fraction (sensitivity) vs true negative fraction (specificity) for different cut-off values between the “true” and “shuffled” distributions. Area under the curve (AUC) reflects the separability between the two distributions. An AUC value close to 1 (100%) indicates that the two distributions are highly separable, which means that the participants on average encoded the temporal measure. The chance level marked as a black circle (and as a vertical dotted line in the density plot) was defined as the cut-off value between the true and the shuffled distribution that showed the most optimal sum of sensitivity (true positive fraction) and specificity (true negative fraction). **a**, Metric time_M1_ involved the degree to which the participant’s response displayed the accurate relative timing of the events or episodes (Temporal pattern_M1_) and accurate timing of the events or episodes relative to the temporal boundaries (Temporal boundary_M1_) (see Figure 1 and Figure S1). Recall accuracy of metric time and chronological time was assessed both within episodes (top row) and between episodes (bottom row). **b**, Object recognition reflects the participants ability to recognize the objects from the object sequences and object association the participants ability to correctly group the objects from the same object sequence (see Figure S1 and Methods). **c**, Metric time_M2_ for four consecutive episodes (1-4, 5-8, 9-12, and 13-16) within each run. Analyses using five consecutive episodes (1-5, 6-10, and 12-16) gave similar results. *P < 0.05 (FDR corrected).

**Figure S3.**
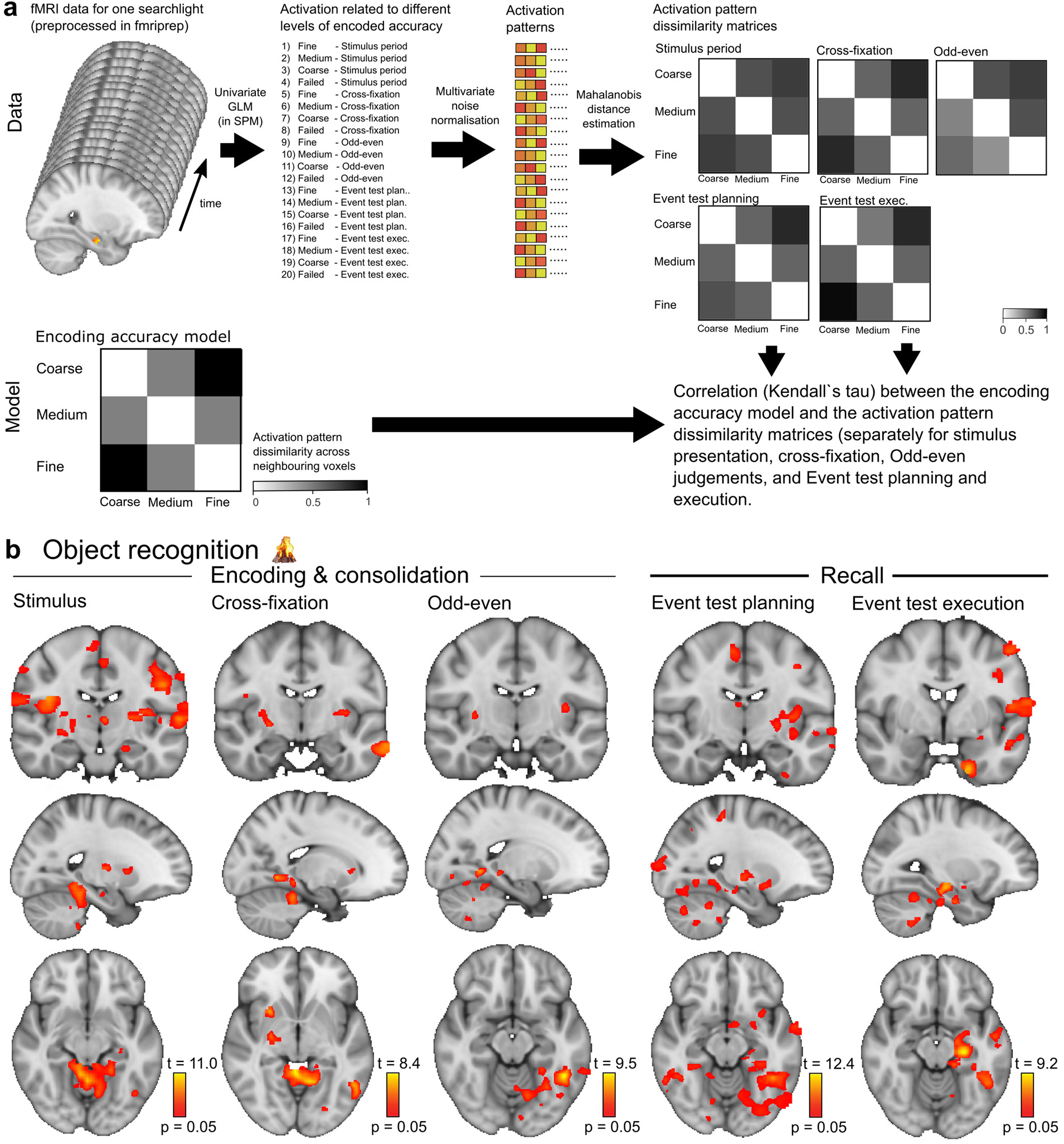
Representation of object identity. **a**, The activation from each searchlight in the brain was first analyzed voxel-wise using a univariate GLM, followed by multivariate noise normalization of the activation patterns related to different levels of encoded accuracy. Activation pattern dissimilarity matrices were then constructed by estimating the Mahalanobis distances between the activation patterns. Finally, the activation pattern dissimilarity matrices were correlated with a model representing consistent modulation of activation pattern dissimilarity with increasing level of encoded accuracy. **b**, Voxels in the brain that showed consistent modulation of activation pattern dissimilarity as object recognition representation became more accurate. Results are shown for the Stimulus period (left), cross-fixation period (mid-left), Odd-even task (mid), and the Event test planning (mid-right) and Event test execution period (right). p = 0.05 represents the cluster mass corrected thresholds.

**Table S1.** Shuffled vs true distributions for temporal and non-temporal measures.

|  | 2-Wasserstein distance<sup>2</sup> | p-value | Specific distribution differences (%) |  |  |
| --- | --- | --- | --- | --- | --- |
|  |  |  | Location | Size | Shape |
| Metric time<sub>M1</sub> |  |  |  |  |  |
| Temporal pattern<sub>M1</sub> | 2.9* | p < 1e-10 | 99 | 0.47 | 0.2 |
| Temporal boundary<sub>M1</sub> | 1.1* | p < 1e-10 | 99 | 0.38 | 1.0 |
| Metric time<sub>M2</sub> |  |  |  |  |  |
| Temporal pattern<sub>M2</sub> | 0.13 | 0.16 | 94 | 0.16 | 6.1 |
| Temporal boundary<sub>M2</sub> | 0.01 | 0.99 | 52 | 1.5 | 46 |
| Chronological time |  |  |  |  |  |
| Event order<sub>C1</sub> | 3.4* | p < 1e-10 | 91 | 8.6 | 0.2 |
| Episode order<sub>C2</sub> | 3.5* | p < 1e-10 | 93 | 7.2 | 0.3 |
| Non-temporal measures |  |  |  |  |  |
| Object recognition | 3.6* | p < 1e-10 | 98 | 1.5 | 0.5 |
| Object association | 2.3* | p < 1e-10 | 66 | 32 | 1.6 |

For each measure, overlap between the true and shuffled distribution was evaluated by using the 2-Wasserstein distance and 5000 bootstrapped samples combined with a generalized Pareto distribution approximation to get accurate p-values. In addition to the squared 2-Wasserstein distance, we also report how much the location, size, and shape of the distributions (in %) contributed to the observed squared 2-Wasserstein distance. \* P < 0.05, corrected for multiple comparisons using a 5% False Discovery Rate (FDR).

**Table S2.** The relationship between temporal and non-temporal measures.

| *Predictors* | **Metric Time<sub>M1</sub>** |  | **Temporal pattern<sub>M2</sub>** |  | **Event order<sub>C1</sub>** |  | **Episode order<sub>C2</sub>** |  | **Object recognition** |  | **Object association** |  |
| --- | --- | --- | --- | --- | --- | --- | --- | --- | --- | --- | --- | --- |
|  | *Estimate* | *Statistic* | *Estimate* | *Statistic* | *Estimate* | *Statistic* | *Estimate* | *Statistic* | *Estimate* | *Statistic* | *Estimate* | *Statistic* |
| (Intercept) | -5.0 | -23 | -5.6 | -2.2 | 4.5 | 10.0 | 1.7 | 5.8 | 3.2 | 5.4 | -0.1 | -0.8 |
| Stimulus period duration | 0.1 | 13.7 |  |  |  |  |  |  |  |  |  |  |
| Q10: I created subgroups of objects | -0.03 | **-2.5\*** | -0.2 | -2.2 |  |  |  |  |  |  |  |  |
| Object sequence number | -0.003 | -3.1 | -0.03 | -6.1 | -0.007 | -4.9 |  |  | 0.011 | 10.9 | 0.003 | 7.4 |
| Group | 0.1 | 2.0 |  |  | -0.6 | -4.3 |  |  | 0.3 | 2.3 |  |  |
| Event test duration | 0.005 | 4.4 | -0.02 | -2.7 | -0.013 | -7.5 |  |  | 0.005 | 4.0 | 0.0001 | 2.4 |
| Q1: I used rhythm | 0.02 | 1.6 | 0.2 | 1.7 |  |  |  |  |  |  |  |  |
| Q21: I mind wandered during odd even | -0.02 | -1.5 |  |  |  |  |  |  | 0.04 | 1.4 |  |  |
| Q17: I mind wandered during cross fix | 0.02 | 1.3 | -0.3 | -2.2 | -0.1 | -2.0 |  |  | -0.1 | **-2.3\*** |  |  |
| **Event order<sub>C1</sub>** | 0.05 | **5.2\*** |  |  |  |  | 0.1 | **4.2\*** | 0.1 | **12.7\*** | 0.011 | **2.4\*** |
| **Episode order<sub>C2</sub>** | -0.006 | -1.1 | 0.1 | **2.4\*** | 0.03 | **3.4\*** |  |  |  |  | 0.011 | **4.0\*** |
| Q2: I counted nr of sec between obj |  |  | 0.3 | **3.2\*** |  |  |  |  |  |  | 0.011 | 1.9 |
| Age |  |  | 0.2 | 3.1 |  |  |  |  | -0.04 | -2.1 |  |  |
| Q9: I imagined moving through an env |  |  | 0.4 | **3.5\*** | 0.1 | **3.8\*** | 0.1 | **2.8\*** | 0.1 | **2.9\*** |  |  |
| Q19: I learned obj seq during odd even |  |  | -0.2 | -1.5 |  |  |  |  |  |  | -0.011 | -1.6 |
| Q4: I gave the objects spatial pos |  |  | -0.2 | -1.6 |  |  |  |  | -0.05 | -1.9 |  |  |
| Q12: I learned obj seq during cross fix |  |  | -0.1 | -1.1 |  |  |  |  | -0.1 | -2.0 |  |  |
| Q6: I thought about obj categories |  |  | 0.1 | 1.3 |  |  |  |  | 0.1 | **2.8\*** |  |  |
| Q3: I used memory aids |  |  | -0.1 | -0.9 | 0.1 | 2.2 | 0.03 | 1.1 |  |  |  |  |
| **Object recognition** |  |  |  |  | 0.3 | **13\*** |  |  |  |  | 0.043 | **6.9\*** |
| Q14: I repeated obj names during cross fix |  |  |  |  | 0.1 | **4.0\*** |  |  | 0.1 | **3.5\*** | -0.018 | **-3.0\*** |
| Q20: I focused on odd even during odd even |  |  |  |  | -0.1 | **-3.1\*** | -0.03 | -1.2 |  |  | -0.018 | **-2.9\*** |
| Q15: I compared obj seq during cross fix |  |  |  |  | -0.1 | -1.7 |  |  |  |  | -0.014 | -1.9 |
| Q8: I thought about obj ass |  |  |  |  | -0.04 | -1.7 | 0.04 | 1.8 |  |  |  |  |
| **Object association** |  |  |  |  | 0.1 | 2.1 | 0.4 | **4.5\*** | 0.2 | **6.0\*** |  |  |
| Q13: I replayed object seq during cross fix |  |  |  |  | -0.04 | -1.3 |  |  |  |  |  |  |
| **Metric Time<sub>M1</sub>** |  |  |  |  | 0.1 | **5.5\*** | -0.03 | -1.0 |  |  |  |  |
| Sex |  |  |  |  |  |  | 0.4 | 1.9 | 0.5 | 1.9 |  |  |
| Q16: I thought about stim period during cross fix |  |  |  |  |  |  | 0.03 | 1.4 |  |  | 0.006 | 1.2 |
| **Temporal Pattern<sub>M2</sub>** |  |  |  |  |  |  | 0.02 | **2.8\*** |  |  |  |  |
| Q22: I remember obj details |  |  |  |  |  |  | -0.03 | -1.3 |  |  |  |  |
| Q11: I gave each object a number |  |  |  |  |  |  |  |  | 0.1 | 1.8 | -0.009 | -1.6 |
| Q5: I made up stories |  |  |  |  |  |  |  |  | -0.04 | -1.8 | 0.029 | **7.3\*** |
| Q7: I thought about object names |  |  |  |  |  |  |  |  |  |  | 0.014 | 2.2 |
| **Random Effects** |  |  |  |  |  |  |  |  |  |  |  |  |
| $\sigma^2$ | 0.81 | | 21.39 | | 1.70 | | 5.17 | | 0.85 | | 0.17 | |
| $\tau_{00}$ | 0.07<sub>ld</sub> | | 4.22<sub>ld</sub> | | 0.28<sub>ld</sub> | | 0.13<sub>ld</sub> | | 0.28<sub>ld</sub> | | 0.01<sub>ld</sub> | |
| ICC | 0.08 |  | 0.16 |  | 0.14 |  | 0.02 |  | 0.25 |  | 0.06 |  |
| N | 101<sub>ld</sub> |  | 101<sub>ld</sub> |  | 98<sub>ld</sub> |  | 101<sub>ld</sub> |  | 98<sub>ld</sub> |  | 98<sub>ld</sub> |  |
| Observations | 4814 |  | 4815 |  | 4670 |  | 4831 |  | 4670 |  | 4670 |  |
| Marginal R<sup>2</sup> / Conditional R<sup>2</sup> | 0.060 / 0.131 |  | 0.085 / 0.236 |  | 0.173 / 0.292 |  | 0.031 / 0.055 |  | 0.198 / 0.397 |  | 0.092 / 0.144 |  |

In these analyses, the temporal and non-temporal measures were employed as response variables in separate models, and explanatory variables were selected on the basis of whether their inclusion improved the second-order AIC (Akaike information criteria) value of the model. In addition, we evaluated absolute measures of goodness-of-fit to determine whether the included variables were indeed informative. We also estimated the variation inflation factors for each model in order to evaluate collinearity between the explanatory variables. The number of observations differed across models because our web-based paradigm failed to register a few responses from two of the participants. Further, the Object recognition score from three participants were excluded because their total number of responses was below 50. Importantly, however, analyses with the outliers included produced similar results and did not change any of the conclusions. \* $P < 0.05$ , corrected for multiple comparisons using a 5% False Discovery Rate (FDR).

**Table S3.** Increased activation pattern dissimilarity for Metric time_M1_

| Brain region | Hemisphere | Cluster nr | Cluster size | t-value(max) | X | Y | Z |
| --- | --- | --- | --- | --- | --- | --- | --- |
| **Stimulus** |  |  |  |  |  |  |  |
| Orbitofrontal cortex | L | 80 | 534 | 8.37 | -33.6 | 34.5 | -21 |
| Temporal pole | L | 80 | 534 | 7.09 | -33.6 | 22.6 | -39 |
| Caudate, anterior | L | 80 | 534 | 6.84 | -9.78 | 7.73 | -9 |
| Temporal fusiform cortex | L | 79 | 92 | 8.44 | -33.6 | -4.16 | -51 |
| Middle temporal gyrus | L | 79 | 92 | 6.34 | -66.3 | -10.1 | -24 |
| Inferior temporal gyrus | L | 79 | 92 | 5.93 | -54.4 | -19 | -24 |
| Middle frontal gyrus | L | 78 | 86 | 6.78 | -54.4 | 16.7 | 36 |
| Inferior frontal gyrus | L | 78 | 86 | 5.89 | -42.5 | 19.6 | 21 |
| Superior frontal gyrus | L | 77 | 69 | 6.01 | -21.7 | 1.79 | 57 |
| Middle frontal gyrus | L | 77 | 69 | 5.55 | -30.6 | -4.16 | 60 |
| Entorhinal cortex, posterior, medial | R | 76 | 62 | 7.11 | 22.9 | -22 | -27 |
| Hippocampus, anterior | R | 76 | 62 | 5.76 | 22.9 | -7.13 | -27 |
| Entorhinal cortex, intermediate, medial | R | 76 | 62 | 5.72 | 19.9 | -13.1 | -27 |
| Middle frontal gyrus | R | 75 | 61 | 5.67 | 55.6 | 22.6 | 33 |
| Inferior frontal gyrus | R | 75 | 61 | 5.32 | 43.7 | 16.7 | 24 |
| Frontal medial cortex | L | 74 | 45 | 6.46 | -0.864 | 46.4 | -12 |
| Putamen, anterior | R | 73 | 40 | 6.91 | 28.9 | 10.7 | 3 |
| Frontal pole | L | 72 | 40 | 7.16 | -24.6 | 52.3 | 3 |
| Orbitofrontal cortex | R | 71 | 37 | 5.6 | 43.7 | 25.6 | -9 |
| Primary somatosensory cortex | L | 70 | 36 | 5.4 | -42.5 | -30.9 | 57 |
| Premotor cortex, BA6 | R | 69 | 30 | 5.62 | 5.08 | -16.1 | 57 |
| Supplementary motor cortex | L | 69 | 30 | 5.53 | -6.81 | -10.1 | 54 |
| Cerebellum, VI | R | 68 | 29 | 6.4 | 22.9 | -54.7 | -24 |
| Superior frontal gyrus | R | 67 | 23 | 6.05 | 25.9 | 1.79 | 72 |
| Supplementary motor cortex | R | 66 | 23 | 5.45 | 2.11 | 1.79 | 57 |
| Thalamus, Prefrontal | R | 65 | 18 | 6.01 | 14 | -4.16 | 12 |
| Orbitofrontal cortex | R | 64 | 18 | 5.49 | 22.9 | 16.7 | -24 |
| Temporal pole | R | 64 | 18 | 5.18 | 28.9 | 13.7 | -30 |
| Cerebellum, VI | L | 63 | 18 | 6 | -30.6 | -39.8 | -33 |
| Entorhinal cortex, intermediate, lateral | L | 62 | 18 | 5.97 | -21.7 | -10.1 | -33 |
| Supramarginal gyrus | L | 61 | 17 | 5.43 | -66.3 | -39.8 | 33 |
| Inferior parietal cortex | L | 61 | 17 | 5.38 | -54.4 | -42.8 | 36 |
| Primary somatosensory cortex | L | 60 | 17 | 5.63 | -33.6 | -39.8 | 66 |
| Entorhinal cortex, posterior, lateral | L | 59 | 16 | 5.66 | -30.6 | -22 | -27 |
| Cerebellum, IIV | R | 58 | 15 | 5.72 | 8.06 | -51.7 | -3 |
| Precentral gyrus | L | 57 | 13 | 5.74 | -36.5 | -16.1 | 51 |
| Premotor cortex, BA6 | L | 57 | 13 | 5.42 | -36.5 | -10.1 | 48 |
| Frontal pole | R | 56 | 11 | 5.21 | 8.06 | 55.3 | 9 |
| Thalamus, Prefrontal | L | 55 | 9 | 5.25 | -12.8 | -7.13 | 12 |
| Paracingulate gyrus | R | 54 | 9 | 6.3 | 8.06 | 28.5 | 45 |
| Secondary somatosensory cortex | L | 53 | 9 | 5.66 | -48.4 | -25 | 18 |
| Middle frontal gyrus | R | 52 | 8 | 5.91 | 37.8 | 31.5 | 18 |
| Anterior intraparietal cortex | L | 51 | 7 | 5.39 | -33.6 | -51.7 | 39 |
| Superior parietal cortex | L | 51 | 7 | 5.13 | -30.6 | -54.7 | 45 |
| Frontal pole | L | 50 | 7 | 5.37 | -18.7 | 70.2 | -3 |
| Cerebellum, IIV | L | 49 | 6 | 5.52 | -0.864 | -48.8 | -6 |
| Frontal pole | R | 48 | 6 | 5.49 | 25.9 | 40.4 | -21 |
| Middle frontal gyrus | R | 47 | 5 | 5.29 | 34.8 | 34.5 | 36 |
| Angular gyrus | R | 46 | 5 | 5.32 | 40.8 | -57.7 | 15 |
| Cerebellum, IIV | L | 45 | 5 | 5.27 | -0.864 | -42.8 | -15 |
| Inferior parietal cortex | L | 44 | 5 | 5.95 | -42.5 | -36.9 | 24 |
| Paracingulate gyrus | L | 43 | 4 | 5.4 | -12.8 | 49.4 | 9 |
| Cingulate gyrus, posterior | R | 42 | 4 | 5.44 | 5.08 | -30.9 | 30 |
| Supramarginal gyrus | L | 41 | 4 | 5.36 | -51.4 | -45.8 | 12 |
| Subcallosal cortex | R | 40 | 4 | 5.22 | 8.06 | 7.73 | -9 |
| Superior parietal cortex | L | 39 | 4 | 5.03 | -39.5 | -51.7 | 57 |
| Postcentral gyrus | L | 38 | 4 | 5.15 | -9.78 | -39.8 | 57 |
| Frontal pole | L | 37 | 4 | 5.38 | -15.7 | 46.4 | -24 |
| Primary somatosensory cortex, BA1 | L | 36 | 4 | 5.07 | -63.3 | -7.13 | 27 |
| Inferior frontal gyrus | L | 35 | 3 | 5.12 | -48.4 | 16.7 | 12 |
| Middle frontal gyrus | L | 34 | 3 | 5.04 | -51.4 | 28.5 | 33 |
| Cerebellum, VI | R | 33 | 3 | 5.15 | 37.8 | -42.8 | -36 |
| Temporal fusiform cortex | R | 32 | 3 | 4.95 | 31.8 | -33.9 | -24 |
| Orbitofrontal cortex | R | 31 | 3 | 5.03 | 17 | 25.6 | -24 |
| Subcallosal cortex | R | 30 | 3 | 5.27 | 8.06 | 19.6 | -24 |
| **Cross-fixation** |  |  |  |  |  |  |  |
| Entorhinal cortex, posterior, medial | R | 5 | 4 | 5.06 | 17 | -19 | -24 |
| **Odd-even** |  |  |  |  |  |  |  |
| na |  |  |  |  |  |  |  |
| **Event test planning** |  |  |  |  |  |  |  |
| Temporal pole | L | 5 | 6 | 5.31 | -27.6 | 19.6 | -39 |
| Entorhinal cortex, posterior, medial | R | 4 | 6 | 5.37 | 17 | -19 | -24 |
| Temporal pole | R | 3 | 3 | 5.43 | 19.9 | 13.7 | -33 |
| Cerebellum, IIV | L | 2 | 3 | 5.04 | -3.84 | -48.8 | -15 |
| **Event test execution** |  |  |  |  |  |  |  |
| Temporal pole | L | 2 | 8 | 6.81 | -24.6 | 19.6 | -39 |
| Cerebellum, IIV | L | 1 | 3 | 5.33 | -0.864 | -48.8 | -6 |

The activation pattern dissimilarity analysis tested whether the activation patterns became more or less similar with increasing encoded accuracy. The activation pattern dissimilarity analysis was carried out using a corrected cluster mass threshold of $p = 0.05$ . Only clusters that were larger than 2 voxels in 2 mm MNI (Montreal Neurological Institute) space were reported, and up to
five local maxima were reported for each cluster. R, right; L, left. X, Y, and Z indicates the position of the max t-value in 2 mm MNI space.

**Table S4.**
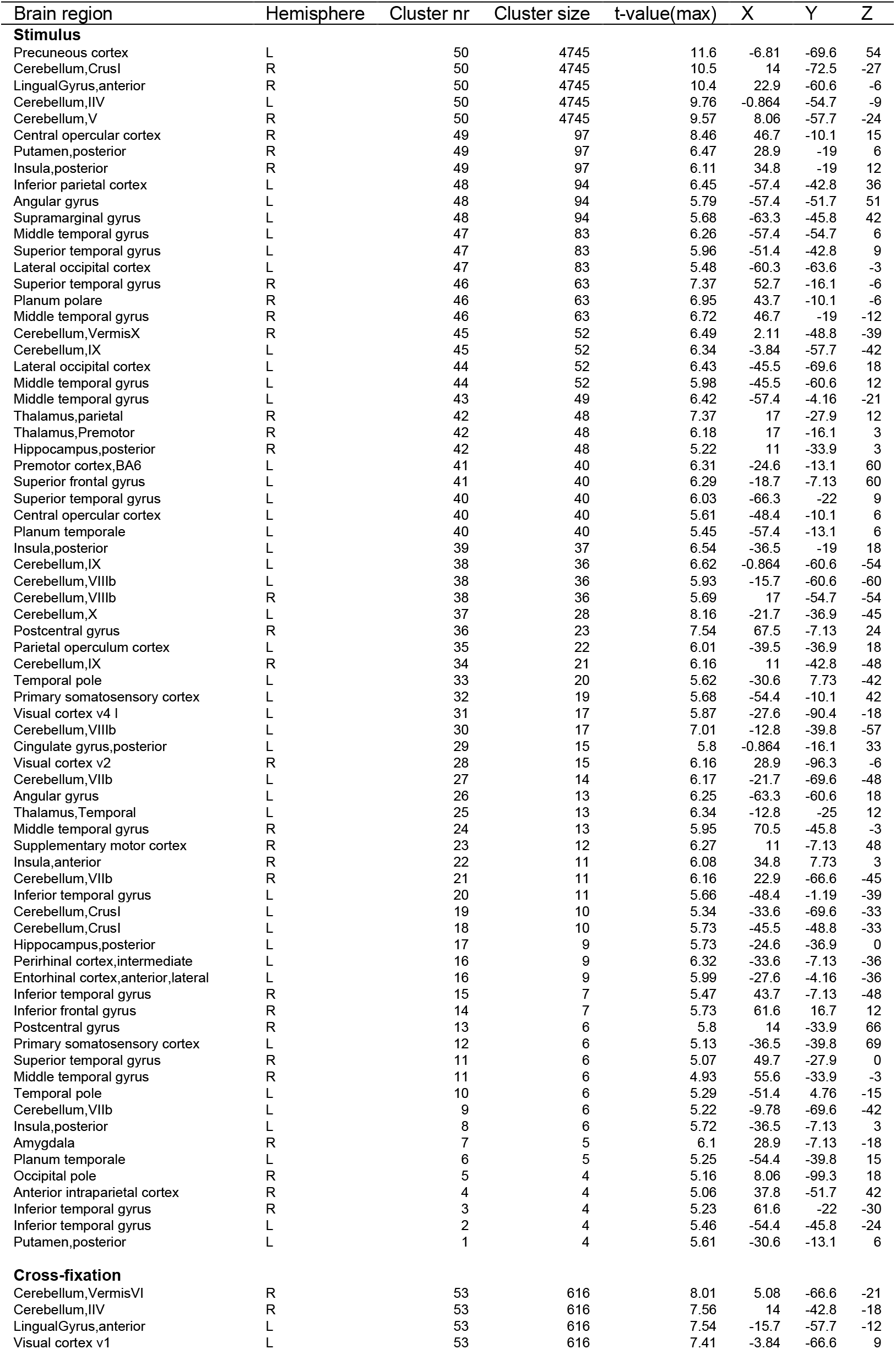
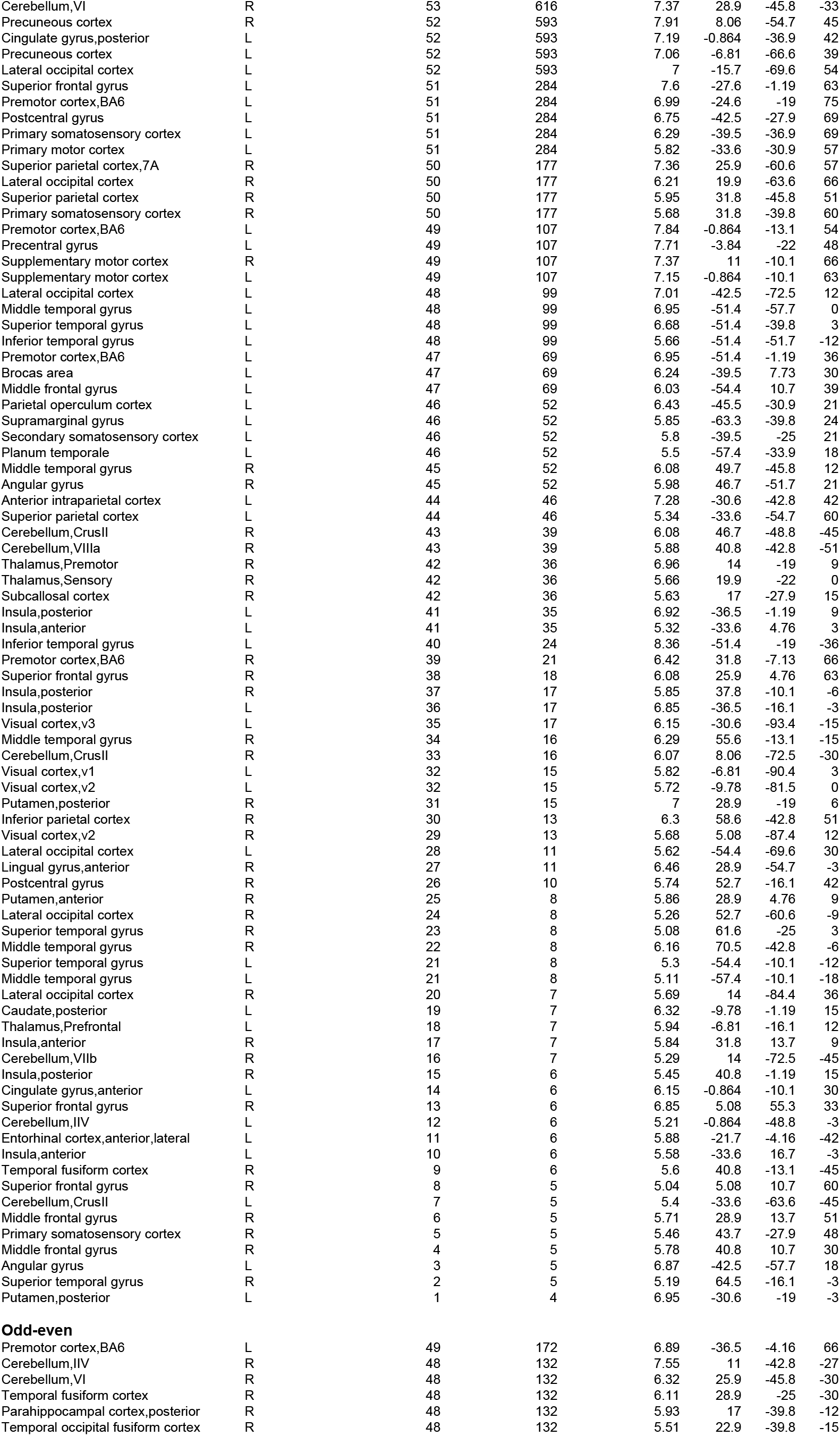
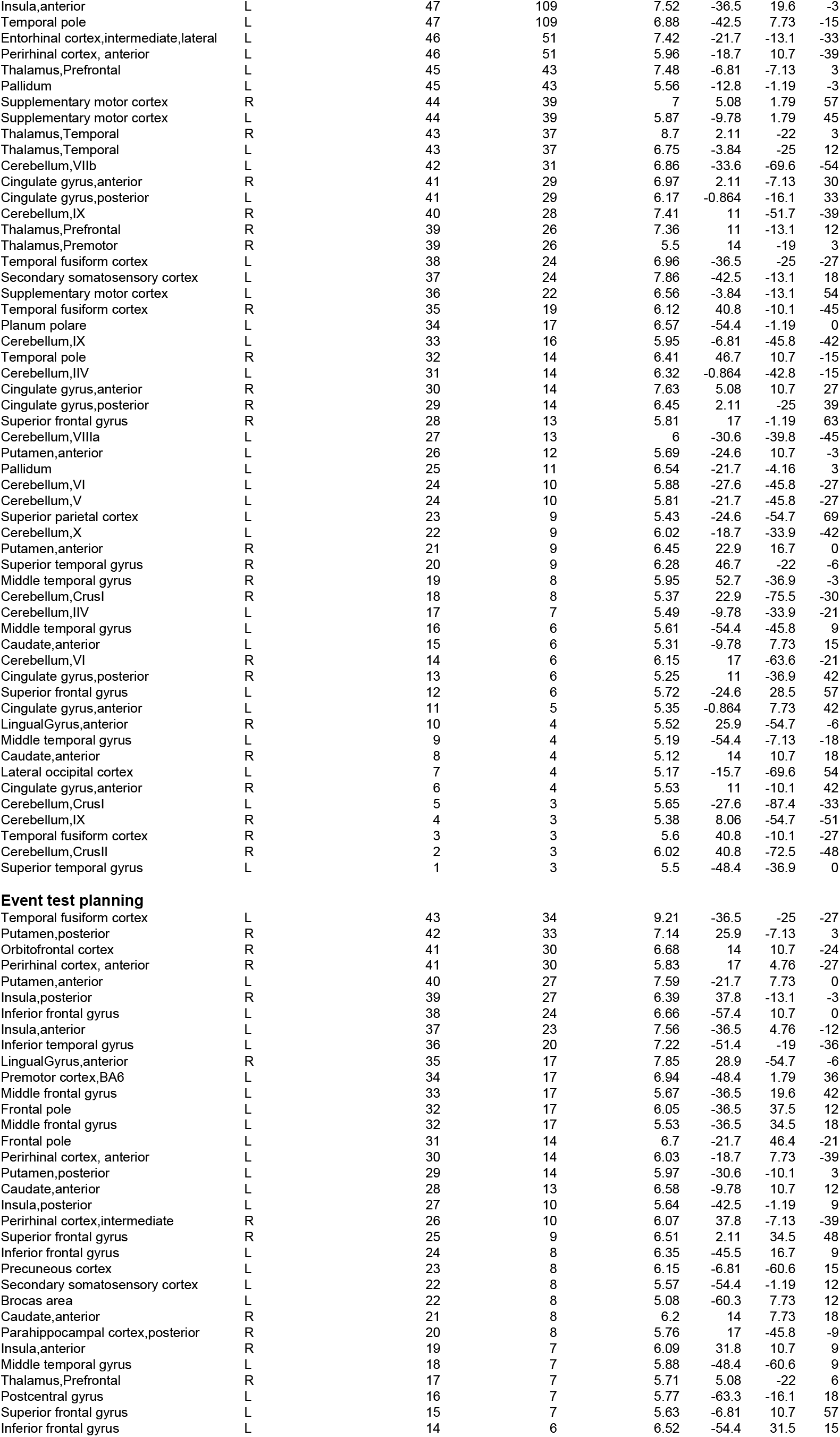
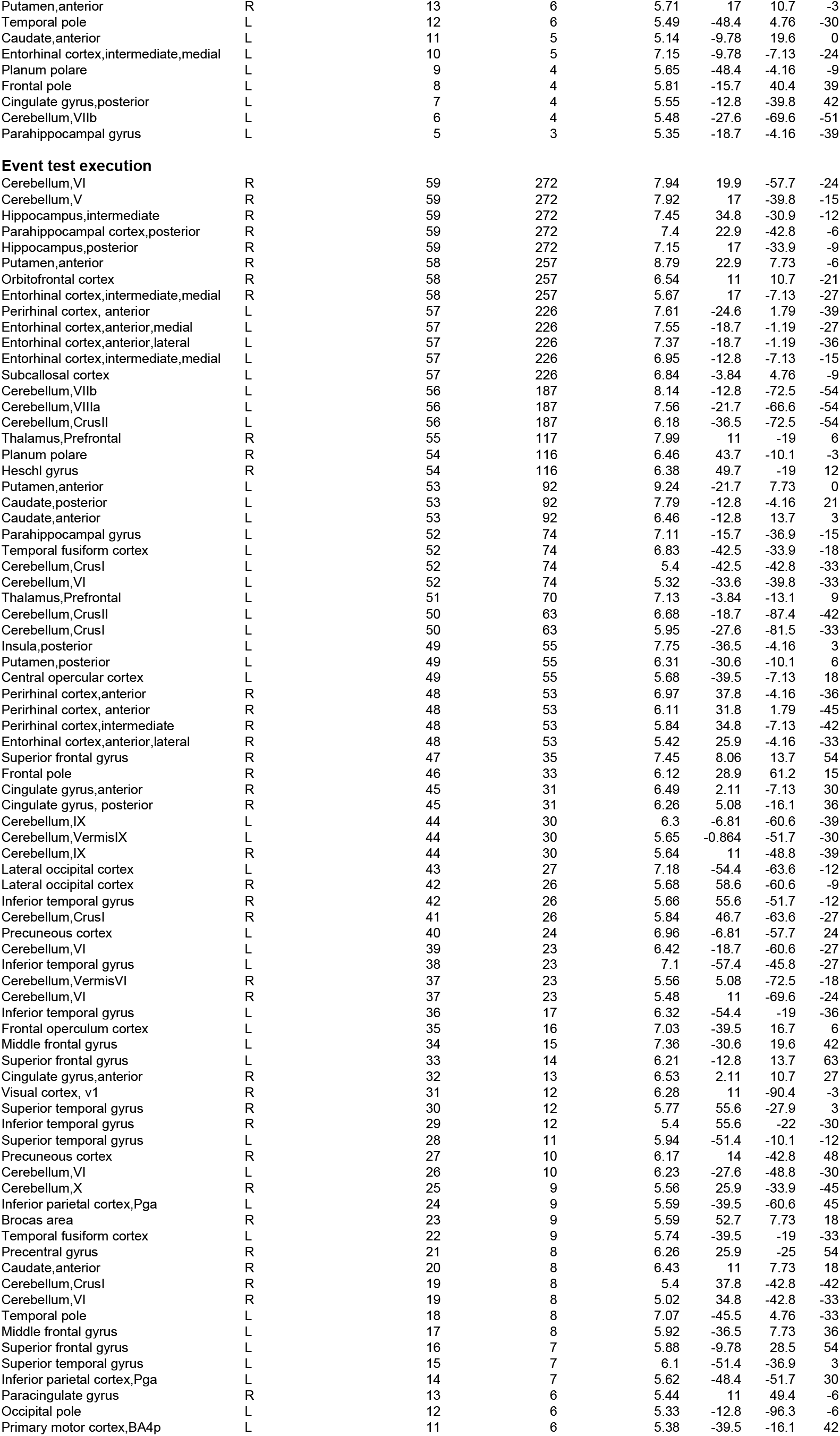
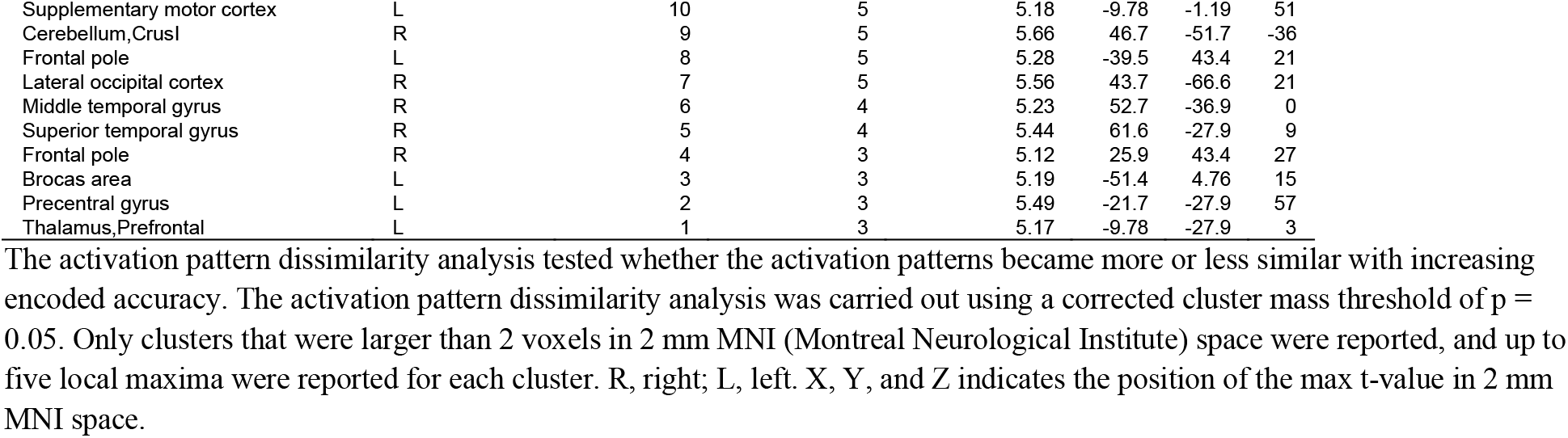
Increased activation pattern dissimilarity for Event order_C1_

**Table S5.**
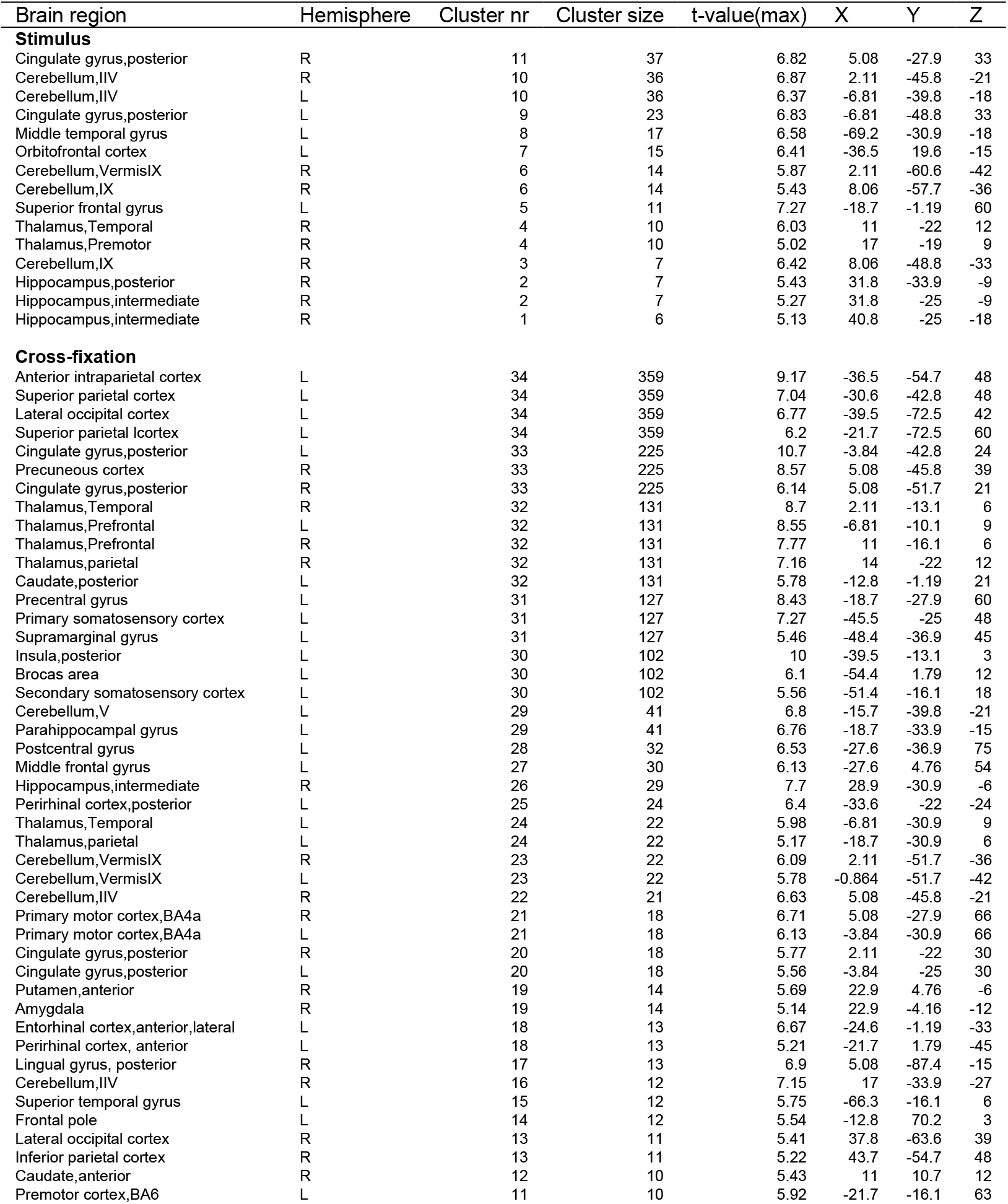
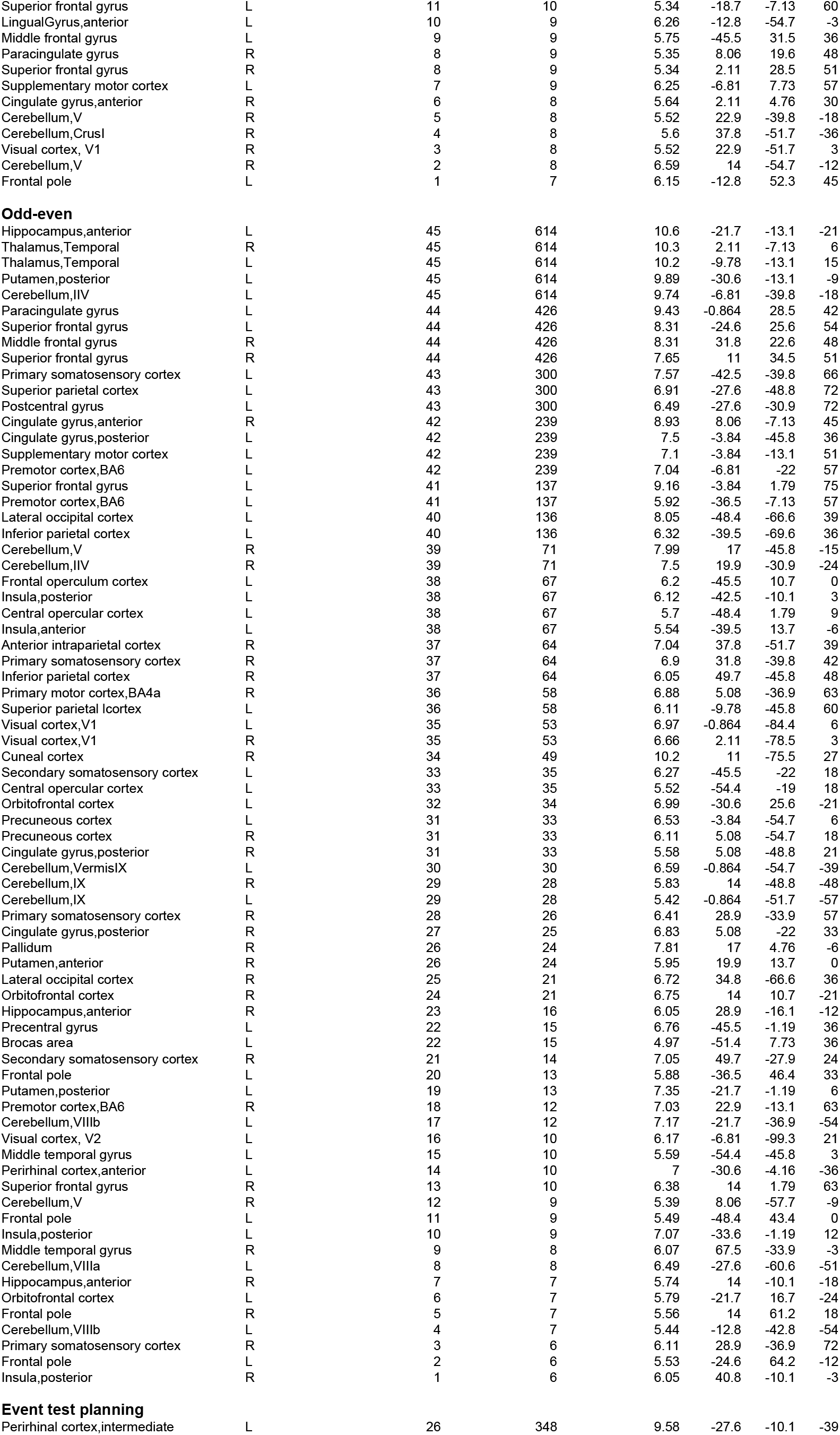
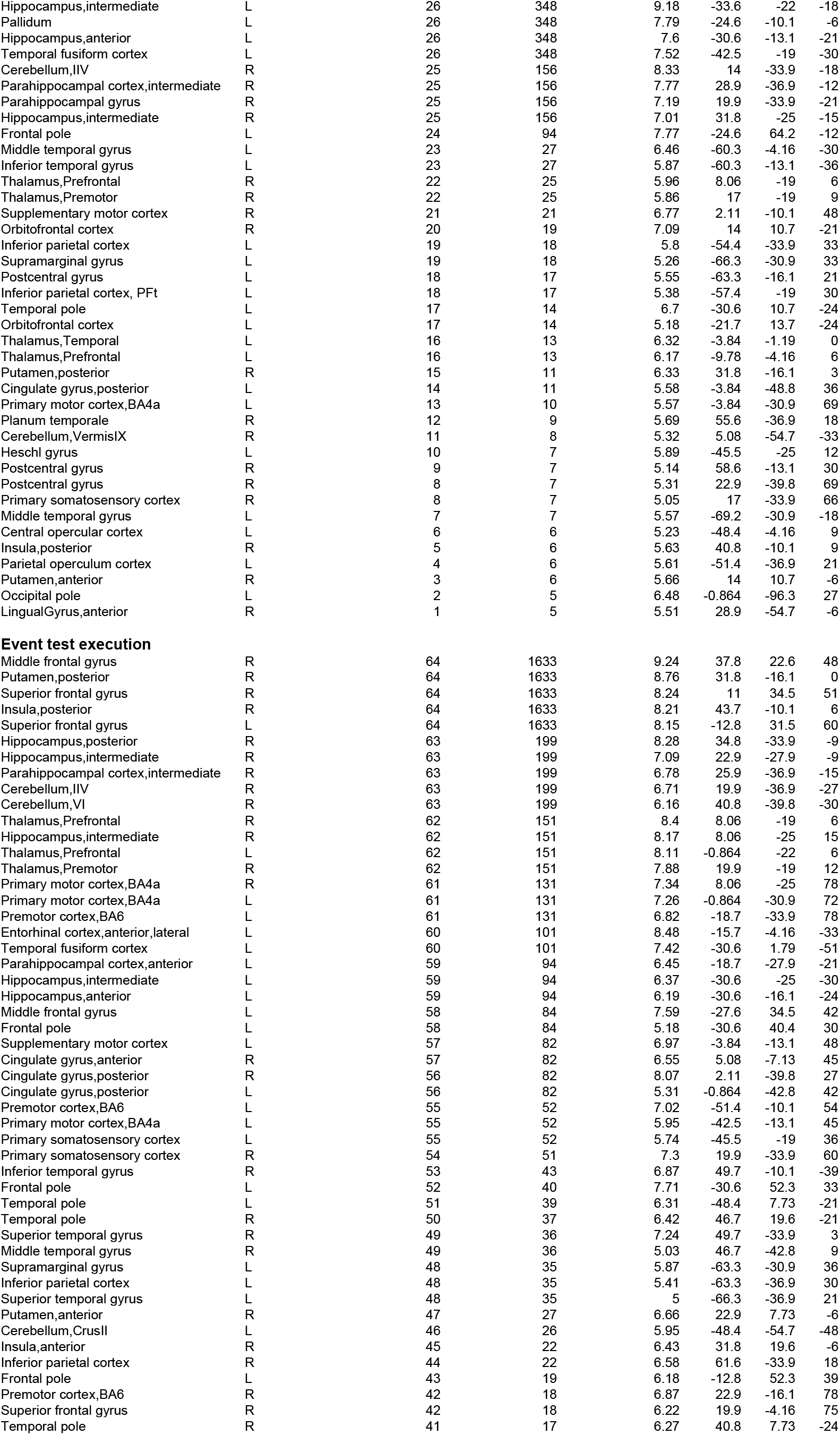
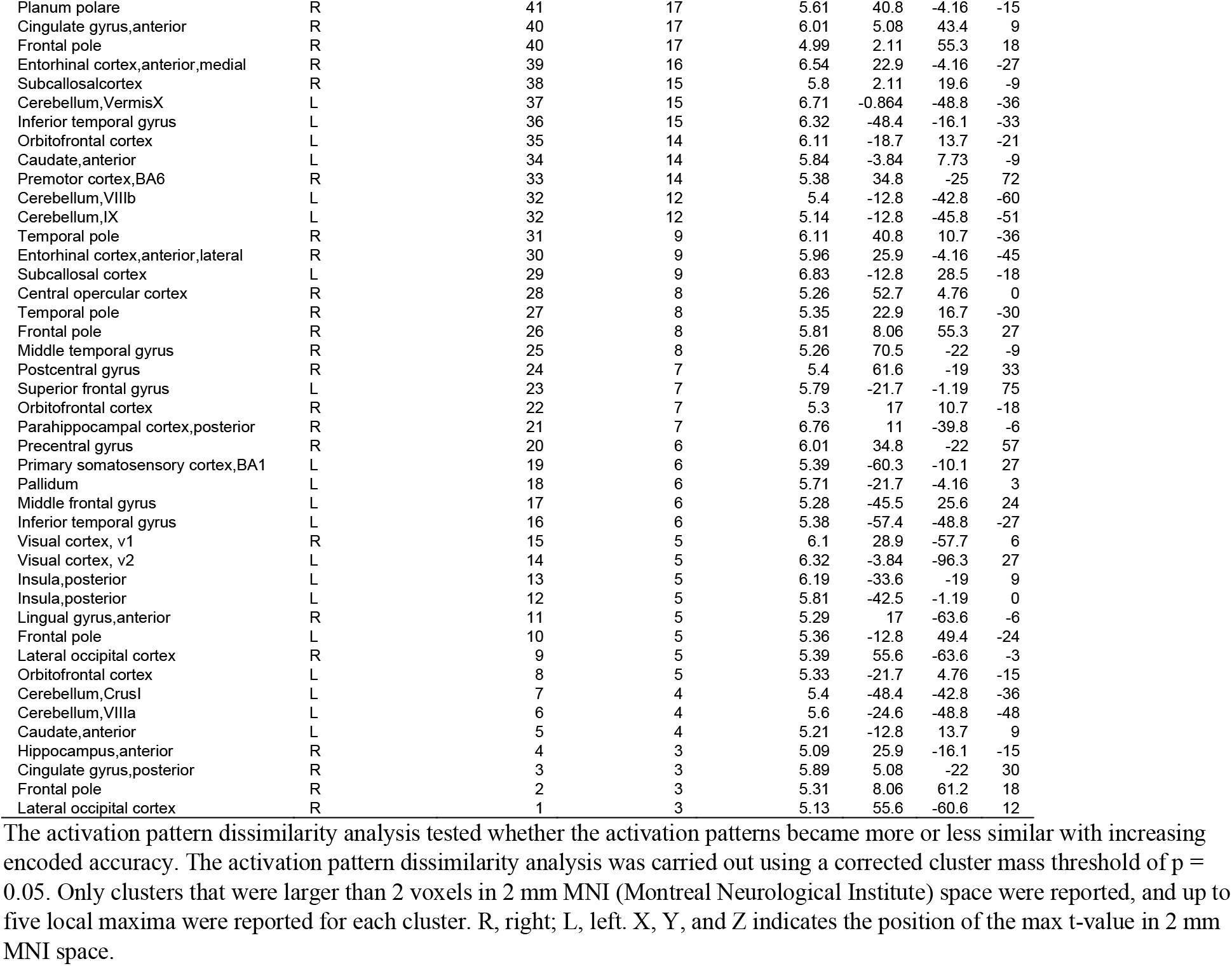
Increased activation pattern dissimilarity for Chronological time_C3_

| Brain region | Hemisphere | Cluster nr | Cluster size | t-value(max) | X | Y | Z |
| --- | --- | --- | --- | --- | --- | --- | --- |
| **Stimulus** |  |  |  |  |  |  |  |
| Cingulate gyrus,posterior | R | 11 | 37 | 6.82 | 5.08 | -27.9 | 33 |
| Cerebellum,IIV | R | 10 | 36 | 6.87 | 2.11 | -45.8 | -21 |
| Cerebellum,IIV | L | 10 | 36 | 6.37 | -6.81 | -39.8 | -18 |
| Cingulate gyrus,posterior | L | 9 | 23 | 6.83 | -6.81 | -48.8 | 33 |
| Middle temporal gyrus | L | 8 | 17 | 6.58 | -69.2 | -30.9 | -18 |
| Orbitofrontal cortex | L | 7 | 15 | 6.41 | -36.5 | 19.6 | -15 |
| Cerebellum,VermisIX | R | 6 | 14 | 5.87 | 2.11 | -60.6 | -42 |
| Cerebellum,IX | R | 6 | 14 | 5.43 | 8.06 | -57.7 | -36 |
| Superior frontal gyrus | L | 5 | 11 | 7.27 | -18.7 | -1.19 | 60 |
| Thalamus,Temporal | R | 4 | 10 | 6.03 | 11 | -22 | 12 |
| Thalamus,Premotor | R | 4 | 10 | 5.02 | 17 | -19 | 9 |
| Cerebellum,IX | R | 3 | 7 | 6.42 | 8.06 | -48.8 | -33 |
| Hippocampus,posterior | R | 2 | 7 | 5.43 | 31.8 | -33.9 | -9 |
| Hippocampus,intermediate | R | 2 | 7 | 5.27 | 31.8 | -25 | -9 |
| Hippocampus,intermediate | R | 1 | 6 | 5.13 | 40.8 | -25 | -18 |
| **Cross-fixation** |  |  |  |  |  |  |  |
| Anterior intraparietal cortex | L | 34 | 359 | 9.17 | -36.5 | -54.7 | 48 |
| Superior parietal cortex | L | 34 | 359 | 7.04 | -30.6 | -42.8 | 48 |
| Lateral occipital cortex | L | 34 | 359 | 6.77 | -39.5 | -72.5 | 42 |
| Superior parietal cortex | L | 34 | 359 | 6.2 | -21.7 | -72.5 | 60 |
| Cingulate gyrus,posterior | L | 33 | 225 | 10.7 | -3.84 | -42.8 | 24 |
| Precuneous cortex | R | 33 | 225 | 8.57 | 5.08 | -45.8 | 39 |
| Cingulate gyrus,posterior | R | 33 | 225 | 6.14 | 5.08 | -51.7 | 21 |
| Thalamus,Temporal | R | 32 | 131 | 8.7 | 2.11 | -13.1 | 6 |
| Thalamus,Prefrontal | L | 32 | 131 | 8.55 | -6.81 | -10.1 | 9 |
| Thalamus,Prefrontal | R | 32 | 131 | 7.77 | 11 | -16.1 | 6 |
| Thalamus,parietal | R | 32 | 131 | 7.16 | 14 | -22 | 12 |
| Caudate,posterior | L | 32 | 131 | 5.78 | -12.8 | -1.19 | 21 |
| Precentral gyrus | L | 31 | 127 | 8.43 | -18.7 | -27.9 | 60 |
| Primary somatosensory cortex | L | 31 | 127 | 7.27 | -45.5 | -25 | 48 |
| Supramarginal gyrus | L | 31 | 127 | 5.46 | -48.4 | -36.9 | 45 |
| Insula,posterior | L | 30 | 102 | 10 | -39.5 | -13.1 | 3 |
| Brocas area | L | 30 | 102 | 6.1 | -54.4 | 1.79 | 12 |
| Secondary somatosensory cortex | L | 30 | 102 | 5.56 | -51.4 | -16.1 | 18 |
| Cerebellum,V | L | 29 | 41 | 6.8 | -15.7 | -39.8 | -21 |
| Parahippocampal gyrus | L | 29 | 41 | 6.76 | -18.7 | -33.9 | -15 |
| Postcentral gyrus | L | 28 | 32 | 6.53 | -27.6 | -36.9 | 75 |
| Middle frontal gyrus | L | 27 | 30 | 6.13 | -27.6 | 4.76 | 54 |
| Hippocampus,intermediate | R | 26 | 29 | 7.7 | 28.9 | -30.9 | -6 |
| Perirhinal cortex,posterior | L | 25 | 24 | 6.4 | -33.6 | -22 | -24 |
| Thalamus,Temporal | L | 24 | 22 | 5.98 | -6.81 | -30.9 | 9 |
| Thalamus,parietal | L | 24 | 22 | 5.17 | -18.7 | -30.9 | 6 |
| Cerebellum,VermisIX | R | 23 | 22 | 6.09 | 2.11 | -51.7 | -36 |
| Cerebellum,VermisIX | L | 23 | 22 | 5.78 | -0.864 | -51.7 | -42 |
| Cerebellum,IIV | R | 22 | 21 | 6.63 | 5.08 | -45.8 | -21 |
| Primary motor cortex,BA4a | R | 21 | 18 | 6.71 | 5.08 | -27.9 | 66 |
| Primary motor cortex,BA4a | L | 21 | 18 | 6.13 | -3.84 | -30.9 | 66 |
| Cingulate gyrus,posterior | R | 20 | 18 | 5.77 | 2.11 | -22 | 30 |
| Cingulate gyrus,posterior | L | 20 | 18 | 5.56 | -3.84 | -25 | 30 |
| Putamen,anterior | R | 19 | 14 | 5.69 | 22.9 | 4.76 | -6 |
| Amygdala | R | 19 | 14 | 5.14 | 22.9 | -4.16 | -12 |
| Entorhinal cortex,anterior,lateral | L | 18 | 13 | 6.67 | -24.6 | -1.19 | -33 |
| Perirhinal cortex, anterior | L | 18 | 13 | 5.21 | -21.7 | 1.79 | -45 |
| Lingual gyrus, posterior | R | 17 | 13 | 6.9 | 5.08 | -87.4 | -15 |
| Cerebellum,IIV | R | 16 | 12 | 7.15 | 17 | -33.9 | -27 |
| Superior temporal gyrus | L | 15 | 12 | 5.75 | -66.3 | -16.1 | 6 |
| Frontal pole | L | 14 | 12 | 5.54 | -12.8 | 70.2 | 3 |
| Lateral occipital cortex | R | 13 | 11 | 5.41 | 37.8 | -63.6 | 39 |
| Inferior parietal cortex | R | 13 | 11 | 5.22 | 43.7 | -54.7 | 48 |
| Caudate,anterior | R | 12 | 10 | 5.43 | 11 | 10.7 | 12 |
| Premotor cortex,BA6 | L | 11 | 10 | 5.92 | -21.7 | -16.1 | 63 |
| Superior frontal gyrus | L | 11 | 10 | 5.34 | -18.7 | -7.13 | 60 |
| LingualGyrus,anterior | L | 10 | 9 | 6.26 | -12.8 | -54.7 | -3 |
| Middle frontal gyrus | L | 9 | 9 | 5.75 | -45.5 | 31.5 | 36 |
| Paracingulate gyrus | R | 8 | 9 | 5.35 | 8.06 | 19.6 | 48 |
| Superior frontal gyrus | R | 8 | 9 | 5.34 | 2.11 | 28.5 | 51 |
| Supplementary motor cortex | L | 7 | 9 | 6.25 | -6.81 | 7.73 | 57 |
| Cingulate gyrus,anterior | R | 6 | 8 | 5.64 | 2.11 | 4.76 | 30 |
| Cerebellum,V | R | 5 | 8 | 5.52 | 22.9 | -39.8 | -18 |
| Cerebellum,CrusI | R | 4 | 8 | 5.6 | 37.8 | -51.7 | -36 |
| Visual cortex, V1 | R | 3 | 8 | 5.52 | 22.9 | -51.7 | 3 |
| Cerebellum,V | R | 2 | 8 | 6.59 | 14 | -54.7 | -12 |
| Frontal pole | L | 1 | 7 | 6.15 | -12.8 | 52.3 | 45 |
| **Odd-even** |  |  |  |  |  |  |  |
| Hippocampus,anterior | L | 45 | 614 | 10.6 | -21.7 | -13.1 | -21 |
| Thalamus,Temporal | R | 45 | 614 | 10.3 | 2.11 | -7.13 | 6 |
| Thalamus,Temporal | L | 45 | 614 | 10.2 | -9.78 | -13.1 | 15 |
| Putamen,posterior | L | 45 | 614 | 9.89 | -30.6 | -13.1 | -9 |
| Cerebellum,IIV | L | 45 | 614 | 9.74 | -6.81 | -39.8 | -18 |
| Paracingulate gyrus | L | 44 | 426 | 9.43 | -0.864 | 28.5 | 42 |
| Superior frontal gyrus | L | 44 | 426 | 8.31 | -24.6 | 25.6 | 54 |
| Middle frontal gyrus | R | 44 | 426 | 8.31 | 31.8 | 22.6 | 48 |
| Superior frontal gyrus | R | 44 | 426 | 7.65 | 11 | 34.5 | 51 |
| Primary somatosensory cortex | L | 43 | 300 | 7.57 | -42.5 | -39.8 | 66 |
| Superior parietal cortex | L | 43 | 300 | 6.91 | -27.6 | -48.8 | 72 |
| Postcentral gyrus | L | 43 | 300 | 6.49 | -27.6 | -30.9 | 72 |
| Cingulate gyrus,anterior | R | 42 | 239 | 8.93 | 8.06 | -7.13 | 45 |
| Cingulate gyrus,posterior | L | 42 | 239 | 7.5 | -3.84 | -45.8 | 36 |
| Supplementary motor cortex | L | 42 | 239 | 7.1 | -3.84 | -13.1 | 51 |
| Premotor cortex,BA6 | L | 42 | 239 | 7.04 | -6.81 | -22 | 57 |
| Superior frontal gyrus | L | 41 | 137 | 9.16 | -3.84 | 1.79 | 75 |
| Premotor cortex,BA6 | L | 41 | 137 | 5.92 | -36.5 | -7.13 | 57 |
| Lateral occipital cortex | L | 40 | 136 | 8.05 | -48.4 | -66.6 | 39 |
| Inferior parietal cortex | L | 40 | 136 | 6.32 | -39.5 | -69.6 | 36 |
| Cerebellum,V | R | 39 | 71 | 7.99 | 17 | -45.8 | -15 |
| Cerebellum,IIV | R | 39 | 71 | 7.5 | 19.9 | -30.9 | -24 |
| Frontal operculum cortex | L | 38 | 67 | 6.2 | -45.5 | 10.7 | 0 |
| Insula,posterior | L | 38 | 67 | 6.12 | -42.5 | -10.1 | 3 |
| Central opercular cortex | L | 38 | 67 | 5.7 | -48.4 | 1.79 | 9 |
| Insula,anterior | L | 38 | 67 | 5.54 | -39.5 | 13.7 | -6 |
| Anterior intraparietal cortex | R | 37 | 64 | 7.04 | 37.8 | -51.7 | 39 |
| Primary somatosensory cortex | R | 37 | 64 | 6.9 | 31.8 | -39.8 | 42 |
| Inferior parietal cortex | R | 37 | 64 | 6.05 | 49.7 | -45.8 | 48 |
| Primary motor cortex,BA4a | R | 36 | 58 | 6.88 | 5.08 | -36.9 | 63 |
| Superior parietal lccortex | L | 36 | 58 | 6.11 | -9.78 | -45.8 | 60 |
| Visual cortex,V1 | L | 35 | 53 | 6.97 | -0.864 | -84.4 | 6 |
| Visual cortex,V1 | R | 35 | 53 | 6.66 | 2.11 | -78.5 | 3 |
| Cuneal cortex | R | 34 | 49 | 10.2 | 11 | -75.5 | 27 |
| Secondary somatosensory cortex | L | 33 | 35 | 6.27 | -45.5 | -22 | 18 |
| Central opercular cortex | L | 33 | 35 | 5.52 | -54.4 | -19 | 18 |
| Orbitofrontal cortex | L | 32 | 34 | 6.99 | -30.6 | 25.6 | -21 |
| Precuneous cortex | L | 31 | 33 | 6.53 | -3.84 | -54.7 | 6 |
| Precuneous cortex | R | 31 | 33 | 6.11 | 5.08 | -54.7 | 18 |
| Cingulate gyrus,posterior | R | 31 | 33 | 5.58 | 5.08 | -48.8 | 21 |
| Cerebellum,VermisIX | L | 30 | 30 | 6.59 | -0.864 | -54.7 | -39 |
| Cerebellum,IX | R | 29 | 28 | 5.83 | 14 | -48.8 | -48 |
| Cerebellum,IX | L | 29 | 28 | 5.42 | -0.864 | -51.7 | -57 |
| Primary somatosensory cortex | R | 28 | 26 | 6.41 | 28.9 | -33.9 | 57 |
| Cingulate gyrus,posterior | R | 27 | 25 | 6.83 | 5.08 | -22 | 33 |
| Pallidum | R | 26 | 24 | 7.81 | 17 | 4.76 | -6 |
| Putamen,anterior | R | 26 | 24 | 5.95 | 19.9 | 13.7 | 0 |
| Lateral occipital cortex | R | 25 | 21 | 6.72 | 34.8 | -66.6 | 36 |
| Orbitofrontal cortex | R | 24 | 21 | 6.75 | 14 | 10.7 | -21 |
| Hippocampus,anterior | R | 23 | 16 | 6.05 | 28.9 | -16.1 | -12 |
| Precentral gyrus | L | 22 | 15 | 6.76 | -45.5 | -1.19 | 36 |
| Brocas area | L | 22 | 15 | 4.97 | -51.4 | 7.73 | 36 |
| Secondary somatosensory cortex | R | 21 | 14 | 7.05 | 49.7 | -27.9 | 24 |
| Frontal pole | L | 20 | 13 | 5.88 | -36.5 | 46.4 | 33 |
| Putamen,posterior | L | 19 | 13 | 7.35 | -21.7 | -1.19 | 6 |
| Premotor cortex,BA6 | R | 18 | 12 | 7.03 | 22.9 | -13.1 | 63 |
| Cerebellum,VIIIb | L | 17 | 12 | 7.17 | -21.7 | -36.9 | -54 |
| Visual cortex, V2 | L | 16 | 10 | 6.17 | -6.81 | -99.3 | 21 |
| Middle temporal gyrus | L | 15 | 10 | 5.59 | -54.4 | -45.8 | 3 |
| Perirhinal cortex,anterior | L | 14 | 10 | 7 | -30.6 | -4.16 | -36 |
| Superior frontal gyrus | R | 13 | 10 | 6.38 | 14 | 1.79 | 63 |
| Cerebellum,V | R | 12 | 9 | 5.39 | 8.06 | -57.7 | -9 |
| Frontal pole | L | 11 | 9 | 5.49 | -48.4 | 43.4 | 0 |
| Insula,posterior | L | 10 | 9 | 7.07 | -33.6 | -1.19 | 12 |
| Middle temporal gyrus | R | 9 | 8 | 6.07 | 67.5 | -33.9 | -3 |
| Cerebellum,VIIla | L | 8 | 8 | 6.49 | -27.6 | -60.6 | -51 |
| Hippocampus,anterior | R | 7 | 7 | 5.74 | 14 | -10.1 | -18 |
| Orbitofrontal cortex | L | 6 | 7 | 5.79 | -21.7 | 16.7 | -24 |
| Frontal pole | R | 5 | 7 | 5.56 | 14 | 61.2 | 18 |
| Cerebellum,VIIIb | L | 4 | 7 | 5.44 | -12.8 | -42.8 | -54 |
| Primary somatosensory cortex | R | 3 | 6 | 6.11 | 28.9 | -36.9 | 72 |
| Frontal pole | L | 2 | 6 | 5.53 | -24.6 | 64.2 | -12 |
| Insula,posterior | R | 1 | 6 | 6.05 | 40.8 | -10.1 | -3 |
| **Event test planning** |  |  |  |  |  |  |  |
| Perirhinal cortex,intermediate | L | 26 | 348 | 9.58 | -27.6 | -10.1 | -39 |
| Hippocampus,intermediate | L | 26 | 348 | 9.18 | -33.6 | -22 | -18 |
| Pallidum | L | 26 | 348 | 7.79 | -24.6 | -10.1 | -6 |
| Hippocampus,anterior | L | 26 | 348 | 7.6 | -30.6 | -13.1 | -21 |
| Temporal fusiform cortex | L | 26 | 348 | 7.52 | -42.5 | -19 | -30 |
| Cerebellum,IIV | R | 25 | 156 | 8.33 | 14 | -33.9 | -18 |
| Parahippocampal cortex,intermediate | R | 25 | 156 | 7.77 | 28.9 | -36.9 | -12 |
| Parahippocampal gyrus | R | 25 | 156 | 7.19 | 19.9 | -33.9 | -21 |
| Hippocampus,intermediate | R | 25 | 156 | 7.01 | 31.8 | -25 | -15 |
| Frontal pole | L | 24 | 94 | 7.77 | -24.6 | 64.2 | -12 |
| Middle temporal gyrus | L | 23 | 27 | 6.46 | -60.3 | -4.16 | -30 |
| Inferior temporal gyrus | L | 23 | 27 | 5.87 | -60.3 | -13.1 | -36 |
| Thalamus,Prefrontal | R | 22 | 25 | 5.96 | 8.06 | -19 | 6 |
| Thalamus,Premotor | R | 22 | 25 | 5.86 | 17 | -19 | 9 |
| Supplementary motor cortex | R | 21 | 21 | 6.77 | 2.11 | -10.1 | 48 |
| Orbitofrontal cortex | R | 20 | 19 | 7.09 | 14 | 10.7 | -21 |
| Inferior parietal cortex | L | 19 | 18 | 5.8 | -54.4 | -33.9 | 33 |
| Supramarginal gyrus | L | 19 | 18 | 5.26 | -66.3 | -30.9 | 33 |
| Postcentral gyrus | L | 18 | 17 | 5.55 | -63.3 | -16.1 | 21 |
| Inferior parietal cortex, PFt | L | 18 | 17 | 5.38 | -57.4 | -19 | 30 |
| Temporal pole | L | 17 | 14 | 6.7 | -30.6 | 10.7 | -24 |
| Orbitofrontal cortex | L | 17 | 14 | 5.18 | -21.7 | 13.7 | -24 |
| Thalamus,Temporal | L | 16 | 13 | 6.32 | -3.84 | -1.19 | 0 |
| Thalamus,Prefrontal | L | 16 | 13 | 6.17 | -9.78 | -4.16 | 6 |
| Putamen,posterior | R | 15 | 11 | 6.33 | 31.8 | -16.1 | 3 |
| Cingulate gyrus,posterior | L | 14 | 11 | 5.58 | -3.84 | -48.8 | 36 |
| Primary motor cortex,BA4a | L | 13 | 10 | 5.57 | -3.84 | -30.9 | 69 |
| Planum temporale | R | 12 | 9 | 5.69 | 55.6 | -36.9 | 18 |
| Cerebellum,VermisIX | R | 11 | 8 | 5.32 | 5.08 | -54.7 | -33 |
| Heschl gyrus | L | 10 | 7 | 5.89 | -45.5 | -25 | 12 |
| Postcentral gyrus | R | 9 | 7 | 5.14 | 58.6 | -13.1 | 30 |
| Postcentral gyrus | R | 8 | 7 | 5.31 | 22.9 | -39.8 | 69 |
| Primary somatosensory cortex | R | 8 | 7 | 5.05 | 17 | -33.9 | 66 |
| Middle temporal gyrus | L | 7 | 7 | 5.57 | -69.2 | -30.9 | -18 |
| Central opercular cortex | L | 6 | 6 | 5.23 | -48.4 | -4.16 | 9 |
| Insula,posterior | R | 5 | 6 | 5.63 | 40.8 | -10.1 | 9 |
| Parietal operculum cortex | L | 4 | 6 | 5.61 | -51.4 | -36.9 | 21 |
| Putamen,anterior | R | 3 | 6 | 5.66 | 14 | 10.7 | -6 |
| Occipital pole | L | 2 | 5 | 6.48 | -0.864 | -96.3 | 27 |
| LingualGyrus,anterior | R | 1 | 5 | 5.51 | 28.9 | -54.7 | -6 |
| **Event test execution** |  |  |  |  |  |  |  |
| Middle frontal gyrus | R | 64 | 1633 | 9.24 | 37.8 | 22.6 | 48 |
| Putamen,posterior | R | 64 | 1633 | 8.76 | 31.8 | -16.1 | 0 |
| Superior frontal gyrus | R | 64 | 1633 | 8.24 | 11 | 34.5 | 51 |
| Insula,posterior | R | 64 | 1633 | 8.21 | 43.7 | -10.1 | 6 |
| Superior frontal gyrus | L | 64 | 1633 | 8.15 | -12.8 | 31.5 | 60 |
| Hippocampus,posterior | R | 63 | 199 | 8.28 | 34.8 | -33.9 | -9 |
| Hippocampus,intermediate | R | 63 | 199 | 7.09 | 22.9 | -27.9 | -9 |
| Parahippocampal cortex,intermediate | R | 63 | 199 | 6.78 | 25.9 | -36.9 | -15 |
| Cerebellum,IIV | R | 63 | 199 | 6.71 | 19.9 | -36.9 | -27 |
| Cerebellum,VI | R | 63 | 199 | 6.16 | 40.8 | -39.8 | -30 |
| Thalamus,Prefrontal | R | 62 | 151 | 8.4 | 8.06 | -19 | 6 |
| Hippocampus,intermediate | R | 62 | 151 | 8.17 | 8.06 | -25 | 15 |
| Thalamus,Prefrontal | L | 62 | 151 | 8.11 | -0.864 | -22 | 6 |
| Thalamus,Premotor | R | 62 | 151 | 7.88 | 19.9 | -19 | 12 |
| Primary motor cortex,BA4a | R | 61 | 131 | 7.34 | 8.06 | -25 | 78 |
| Primary motor cortex,BA4a | L | 61 | 131 | 7.26 | -0.864 | -30.9 | 72 |
| Premotor cortex,BA6 | L | 61 | 131 | 6.82 | -18.7 | -33.9 | 78 |
| Entorhinal cortex,anterior,lateral | L | 60 | 101 | 8.48 | -15.7 | -4.16 | -33 |
| Temporal fusiform cortex | L | 60 | 101 | 7.42 | -30.6 | 1.79 | -51 |
| Parahippocampal cortex,anterior | L | 59 | 94 | 6.45 | -18.7 | -27.9 | -21 |
| Hippocampus,intermediate | L | 59 | 94 | 6.37 | -30.6 | -25 | -30 |
| Hippocampus,anterior | L | 59 | 94 | 6.19 | -30.6 | -16.1 | -24 |
| Middle frontal gyrus | L | 58 | 84 | 7.59 | -27.6 | 34.5 | 42 |
| Frontal pole | L | 58 | 84 | 5.18 | -30.6 | 40.4 | 30 |
| Supplementary motor cortex | L | 57 | 82 | 6.97 | -3.84 | -13.1 | 48 |
| Cingulate gyrus,anterior | R | 57 | 82 | 6.55 | 5.08 | -7.13 | 45 |
| Cingulate gyrus,posterior | R | 56 | 82 | 8.07 | 2.11 | -39.8 | 27 |
| Cingulate gyrus,posterior | L | 56 | 82 | 5.31 | -0.864 | -42.8 | 42 |
| Premotor cortex,BA6 | L | 55 | 52 | 7.02 | -51.4 | -10.1 | 54 |
| Primary motor cortex,BA4a | L | 55 | 52 | 5.95 | -42.5 | -13.1 | 45 |
| Primary somatosensory cortex | L | 55 | 52 | 5.74 | -45.5 | -19 | 36 |
| Primary somatosensory cortex | R | 54 | 51 | 7.3 | 19.9 | -33.9 | 60 |
| Inferior temporal gyrus | R | 53 | 43 | 6.87 | 49.7 | -10.1 | -39 |
| Frontal pole | L | 52 | 40 | 7.71 | -30.6 | 52.3 | 33 |
| Temporal pole | L | 51 | 39 | 6.31 | -48.4 | 7.73 | -21 |
| Temporal pole | R | 50 | 37 | 6.42 | 46.7 | 19.6 | -21 |
| Superior temporal gyrus | R | 49 | 36 | 7.24 | 49.7 | -33.9 | 3 |
| Middle temporal gyrus | R | 49 | 36 | 5.03 | 46.7 | -42.8 | 9 |
| Supramarginal gyrus | L | 48 | 35 | 5.87 | -63.3 | -30.9 | 36 |
| Inferior parietal cortex | L | 48 | 35 | 5.41 | -63.3 | -36.9 | 30 |
| Superior temporal gyrus | L | 48 | 35 | 5 | -66.3 | -36.9 | 21 |
| Putamen,anterior | R | 47 | 27 | 6.66 | 22.9 | 7.73 | -6 |
| Cerebellum,CrusII | L | 46 | 26 | 5.95 | -48.4 | -54.7 | -48 |
| Insula,anterior | R | 45 | 22 | 6.43 | 31.8 | 19.6 | -6 |
| Inferior parietal cortex | R | 44 | 22 | 6.58 | 61.6 | -33.9 | 18 |
| Frontal pole | L | 43 | 19 | 6.18 | -12.8 | 52.3 | 39 |
| Premotor cortex,BA6 | R | 42 | 18 | 6.87 | 22.9 | -16.1 | 78 |
| Superior frontal gyrus | R | 42 | 18 | 6.22 | 19.9 | -4.16 | 75 |
| Temporal pole | R | 41 | 17 | 6.27 | 40.8 | 7.73 | -24 |
| Planum polare | R | 41 | 17 | 5.61 | 40.8 | -4.16 | -15 |
| Cingulate gyrus,anterior | R | 40 | 17 | 6.01 | 5.08 | 43.4 | 9 |
| Frontal pole | R | 40 | 17 | 4.99 | 2.11 | 55.3 | 18 |
| Entorhinal cortex,anterior,medial | R | 39 | 16 | 6.54 | 22.9 | -4.16 | -27 |
| Subcallosalcortex | R | 38 | 15 | 5.8 | 2.11 | 19.6 | -9 |
| Cerebellum,VermisX | L | 37 | 15 | 6.71 | -0.864 | -48.8 | -36 |
| Inferior temporal gyrus | L | 36 | 15 | 6.32 | -48.4 | -16.1 | -33 |
| Orbitofrontal cortex | L | 35 | 14 | 6.11 | -18.7 | 13.7 | -21 |
| Caudate,anterior | L | 34 | 14 | 5.84 | -3.84 | 7.73 | -9 |
| Premotor cortex,BA6 | R | 33 | 14 | 5.38 | 34.8 | -25 | 72 |
| Cerebellum,VIIIb | L | 32 | 12 | 5.4 | -12.8 | -42.8 | -60 |
| Cerebellum,IX | L | 32 | 12 | 5.14 | -12.8 | -45.8 | -51 |
| Temporal pole | R | 31 | 9 | 6.11 | 40.8 | 10.7 | -36 |
| Entorhinal cortex,anterior,lateral | R | 30 | 9 | 5.96 | 25.9 | -4.16 | -45 |
| Subcallosal cortex | L | 29 | 9 | 6.83 | -12.8 | 28.5 | -18 |
| Central opercular cortex | R | 28 | 8 | 5.26 | 52.7 | 4.76 | 0 |
| Temporal pole | R | 27 | 8 | 5.35 | 22.9 | 16.7 | -30 |
| Frontal pole | R | 26 | 8 | 5.81 | 8.06 | 55.3 | 27 |
| Middle temporal gyrus | R | 25 | 8 | 5.26 | 70.5 | -22 | -9 |
| Postcentral gyrus | R | 24 | 7 | 5.4 | 61.6 | -19 | 33 |
| Superior frontal gyrus | L | 23 | 7 | 5.79 | -21.7 | -1.19 | 75 |
| Orbitofrontal cortex | R | 22 | 7 | 5.3 | 17 | 10.7 | -18 |
| Parahippocampal cortex,posterior | R | 21 | 7 | 6.76 | 11 | -39.8 | -6 |
| Precentral gyrus | R | 20 | 6 | 6.01 | 34.8 | -22 | 57 |
| Primary somatosensory cortex,BA1 | L | 19 | 6 | 5.39 | -60.3 | -10.1 | 27 |
| Pallidum | L | 18 | 6 | 5.71 | -21.7 | -4.16 | 3 |
| Middle frontal gyrus | L | 17 | 6 | 5.28 | -45.5 | 25.6 | 24 |
| Inferior temporal gyrus | L | 16 | 6 | 5.38 | -57.4 | -48.8 | -27 |
| Visual cortex, v1 | R | 15 | 5 | 6.1 | 28.9 | -57.7 | 6 |
| Visual cortex, v2 | L | 14 | 5 | 6.32 | -3.84 | -96.3 | 27 |
| Insula,posterior | L | 13 | 5 | 6.19 | -33.6 | -19 | 9 |
| Insula,posterior | L | 12 | 5 | 5.81 | -42.5 | -1.19 | 0 |
| Lingual gyrus,anterior | R | 11 | 5 | 5.29 | 17 | -63.6 | -6 |
| Frontal pole | L | 10 | 5 | 5.36 | -12.8 | 49.4 | -24 |
| Lateral occipital cortex | R | 9 | 5 | 5.39 | 55.6 | -63.6 | -3 |
| Orbitofrontal cortex | L | 8 | 5 | 5.33 | -21.7 | 4.76 | -15 |
| Cerebellum,CrusI | L | 7 | 4 | 5.4 | -48.4 | -42.8 | -36 |
| Cerebellum,VIIla | L | 6 | 4 | 5.6 | -24.6 | -48.8 | -48 |
| Caudate,anterior | L | 5 | 4 | 5.21 | -12.8 | 13.7 | 9 |
| Hippocampus,anterior | R | 4 | 3 | 5.09 | 25.9 | -16.1 | -15 |
| Cingulate gyrus,posterior | R | 3 | 3 | 5.89 | 5.08 | -22 | 30 |
| Frontal pole | R | 2 | 3 | 5.31 | 8.06 | 61.2 | 18 |
| Lateral occipital cortex | R | 1 | 3 | 5.13 | 55.6 | -60.6 | 12 |

The activation pattern dissimilarity analysis tested whether the activation patterns became more or less similar with increasing encoded accuracy. The activation pattern dissimilarity analysis was carried out using a corrected cluster mass threshold of $p = 0.05$ . Only clusters that were larger than 2 voxels in 2 mm MNI (Montreal Neurological Institute) space were reported, and up to five local maxima were reported for each cluster. R, right; L, left. X, Y, and Z indicates the position of the max t-value in 2 mm MNI space.

**Table S6.**
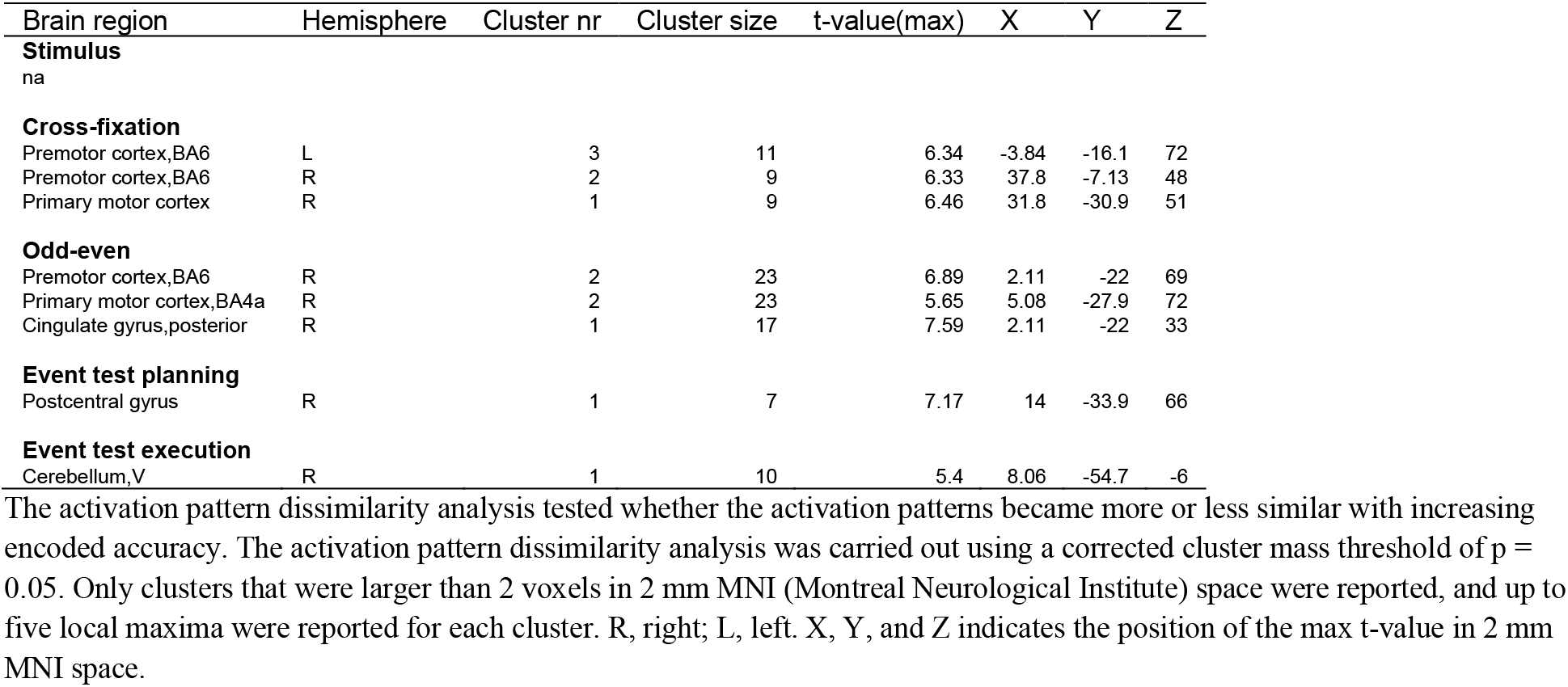
Increased activation pattern dissimilarity for Episode order_C2_

